# Nutrient-regulated dynamics of chondroprogenitors in the postnatal murine growth plate

**DOI:** 10.1101/2023.01.20.524764

**Authors:** Takeshi Oichi, Joe Kodama, Kimberly Wilson, Hongying Tian, Yuka Imamura, Yu Usami, Yasushi Oshima, Taku Saito, Sakae Tanaka, Masahiro Iwamoto, Satoru Otsuru, Motomi Iwamoto-Enomoto

## Abstract

Longitudinal bone growth relies on endochondral ossification in the cartilaginous growth plate where chondrocytes accumulate and synthesize the matrix scaffold that is replaced by bone. The chondroprogenitors in the resting zone maintain the continuous turnover of chondrocytes in the growth plate. Malnutrition is a leading cause of growth retardation in children; however, after recovery from nutrient deprivation, bone growth is accelerated beyond the normal rate, a phenomenon termed catch-up growth. Though nutritional status is a known regulator of long bone growth, it is largely unknown if and how chondroprogenitor cells respond to deviations in nutrient availability. Here, using fate-mapping analysis in *Axin2Cre^ERT2^* mice, we showed that dietary restriction increased the number of Axin2+ chondroprogenitors in the resting zone and simultaneously inhibited their differentiation. Once nutrient deficiency was resolved, the accumulated chondroprogenitor cells immediately restarted differentiation and formed chondrocyte columns, contributing to accelerated growth. Furthermore, we showed that nutrient deprivation reduced the level of phosphorylated Akt in the resting zone, and that exogenous IGF-1 canceled this reduction and stimulated differentiation of the pooled chondroprogenitors, decreasing their numbers. Our study of *Axin2Cre^ERT2^* revealed that nutrient availability regulates the balance between accumulation and differentiation of chondroprogenitors in the growth plate, and further demonstrated that IGF-1 partially mediates this regulation by promoting the committed differentiation of the chondroprogenitor cells.

## INTRODUCTION

Long bone growth is promoted by endochondral ossification that occurs at growth plates, which are cartilaginous tissues at the ends of long bones and vertebral bodies.^1^. The growth plate comprises three layers: the resting, proliferative, and hypertrophic zones. The resting zone contains round, undifferentiated chondrocytes (hereafter “chondroprogenitors”) that rarely divide and are precursors to proliferative chondrocytes^2,3^ Recent lineage-tracing studies have identified chondroprogenitors in the resting zone that contribute continually to long bone growth via the acquisition of stem-cell characteristics after the formation of the secondary ossification center (SOC).^4,5^ The analysis of the genome-wide association study of human height reveals that specificity of gene expression in the resting chondrocytes in newborn mice is significantly associated with height GWAS *p*-values, stressing the important role of these cell populations in determining the human skeletal growth.^6^

Malnutrition is considered a leading cause of growth retardation in children. ^7,8^ When transient nutritional impairment is resolved, long bone growth rate often accelerates beyond the normal rate; this phenomenon is termed catch-up growth.^9^ A common hypothesis to explain catch-up growth is that growth-inhibiting conditions decrease the proliferation of chondroprogenitors, preserving their proliferation potential until the growth-inhibiting conditions are resolved, at which point the preserved proliferation capacity accelerates growth.^10^ Although this hypothesis suggests that chondroprogenitors play an important role in this phenomenon, the effect of nutritional availability on their function is poorly understood. One reason for the limited research on this topic is the lack of appropriate tools to mark and visualize chondroprogenitors at appropriate times.

Wnt proteins participate in the maintenance of stem cells in many adult mammalian tissues ^11^ Wnt signaling is mediated by a pathway that activates transcription factors via the intracellular protein β-catenin. Axin2, a universal transcriptional target of β-catenin-dependent Wnt signaling, provides a reliable readout of cell responses to Wnt.^11,12^ Genetic lineage tracing of the cells marked by the reporter protein expression in the *Axin2Cre^ERT2^* mice has successfully identified stem cells in several adult mammalian tissues.^13–16^ We used this lineage tracing approach to identify a unique population of Axin2+ chondroprogenitor cells that reside in the resting zone of the growth plate after SOC formation. In this study, we investigated the nutrient-regulated dynamics of these chondroprogenitors and their regulatory mechanism using fate mapping analysis with a mouse nutrient-induced catch-up growth model.

## RESULTS

### Identification of Axin2^+^ Cells in the Resting Zone of the Growth Plate after SOC Formation

We first investigated whether Axin2-expressing cells contain chondroprogenitors in the resting zone of the growth plate, because *Axin2* gene has been successfully used to identify tissue-specific stem cells in other tissues.^13–16^ The administration of tamoxifen for three consecutive days induced Cre-mediated expression of the R26RZsGreen reporter, allowing us to observe that Axin2^+^ cells were rarely present in the resting zone of the neonatal growth plate (Fig. 1a and e). However, they appeared in the resting zone after SOC formation (Fig. 1b, proximal tibia and Fig. S1, distal femur), which is consistent with a previous study^17^ and the number of these Axin2^+^ cells in the resting zone decreased when tamoxifen induction was performed at older ages (Fig. 1c–e). The highly specific localization of Axin2^+^ cells in the growth plate was observed at all ages after SOC formation, suggesting that chondroprogenitor cells in the resting zone were labeled.

**Figure 1.**
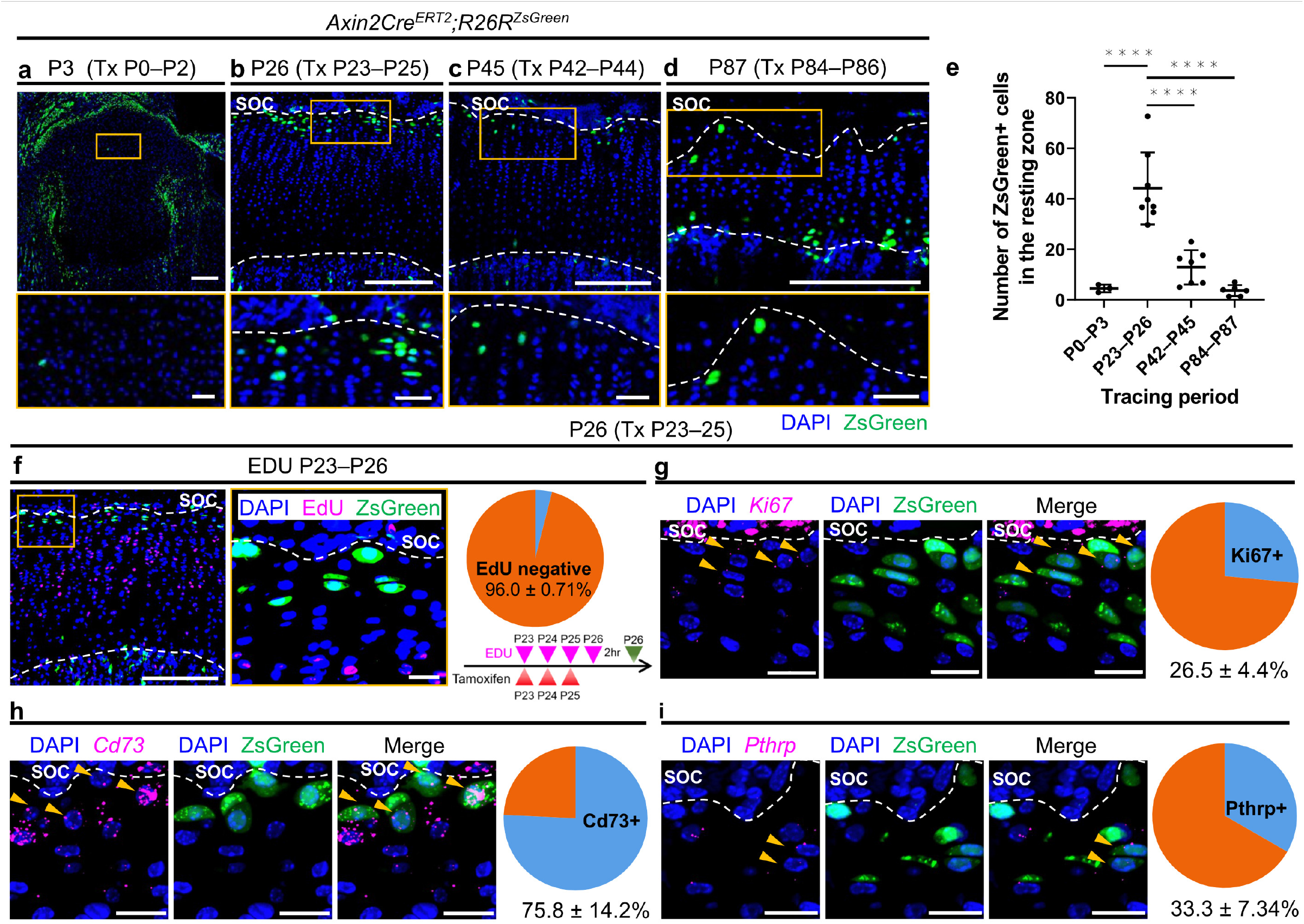
Identification and characterization of Axin2^+^ cells in the resting zone after secondary ossification center formation. **a–d** Axin2+ cells in the proximal tibial growth plate in *Axin2Cre^ERT2^;R26R^ZsGreen^* mice. Axin2^+^ cells were labelled one day after three consecutive daily injections of tamoxifen (120 μg/g body weight) from P0 (n = 3) (**a**), P23 (n = 8) (**b**), P42 (n = 7) (**c**), and P84 (n = 6) (**d**). Bottom panels show magnified views of the resting zone of the growth plate. **e** Quantification of ZsGreen^+^ cells in the resting zone across different time points reported above in (**a–d**). Adjusted ****p= < 0.0001. **f–i** Characterization of Axin2^+^ cells in the resting zone of the proximal tibial growth plate in *Axin2Cre^ERT2^;R26R^ZsGreen^* mice one day after three consecutive daily tamoxifen injections from P23. **f** Representative images of the EdU labeling and Axin2^+^ cells, and percentage of EdU negative cells among Axin2+ cells (n = 3). EdU (30 μg/g body weight) was injected daily from P23 to P26 and mice were euthanized 2 hours after last administration of EdU. **g** Representative images for *in situ* hybridization of *Ki67* and Axin2^+^ cells, and percentage of *Ki67*^+^cells among Axin2^+^ cells (n = 3). Arrow heads, Axin2^+^ cells expressing *Ki67*. **h** Representative images for *in situ* hybridization of *Cd73* and Axin2^+^ cells, and percentage of *Cd73*^+^ cells among Axin2^+^ cells (n = 3). Arrow heads, Axin2^+^ cells expressing *Cd73*. **i** Representative images for *in situ* hybridization of *Pthrp* and Axin2^+^ cells, and percentage of *Pthrp*^+^ cells among Axin2^+^ cells (n = 3). Arrow heads, Axin2^+^ cells expressing *Pthrp*. Tx, Tamoxifen, SOC, secondary ossification center. The white dashed lines demarcate the growth plate from the surrounding tissues. Scale bars: 200 μm (**a–d** [upper panels], **f** [left]), 40 μm (**a–d** [bottom panels]), 20 μm (**f** [right], **g–i**). All data are presented as the mean ± SD. Statistical significance was determined by one-way analysis of variance and Tukey’s multiple comparison test.

We focused on the Axin2^+^ cells in the resting zone at P23 because the SOC is fully developed by this time and the growth plate activity is high at this time.^5^ To characterize these Axin2+ cells, the chondrocytes were isolated from the growth plate of *Axin2Cre^ERT2^;R26R^ZsGreen^* mice at P26 after tamoxifen injections at P23-P25 and were fractionated by skeletal stem cell markers as reported previously (Fig. S2a).^18^ The Axin2+ cells were distributed into various types of stem cell populations, including ‘multipotent progenitors’ and ‘stromal progenitors’ (Fig. S2b), and 41.8% of total Axin2+ cells were fractionated in the ‘multipotent progenitor’ population as characterized by CD45-TER119-Tie2-AlphaV+Thy-6C3-CD105-CD200+ (Fig. S2c). To test whether Axin2+ cells and their descendants have stem cell like properties in cultured conditions, the cells were isolated from the growth plate of *Axin2Cre^ERT2^;R26R^ZsGreen^* mice at P37 after tamoxifen injections at P23-P25 and were expanded by serial passaging. At Passage 6, the cells stopped proliferating: the population doubling level when they ceased proliferation was approximately 13.0. The percentage of ZsGreen-positive cells were 32 ± 7.5% at Passage 3 and 42 ± 8.2% at Passage 5. At Passage 3, the ZsGreen-positive cells were sorted and cultured under chondrogenic, osteogenic and adipogenic differentiation conditions. The ZsGreen-positive cells showed accumulation of sulfated proteoglycan, calcification and oil droplet under chondrogenic, osteogenic and adipogenic differentiation conditions, respectively (Fig. S2d). These results suggest that the Axin2+ cells have chondrogenic, osteogenic and adipogenic differentiation abilities, but their expansion property is limited.

To examine the proliferation activity of these Axin2^+^ cells, we performed an EdU label-exclusive assay and *in situ* hybridization of *Mki67* (*Ki67)*. Most Axin2^+^ cells excluded EdU incorporation (Fig. 1f, EdU^-^; 96.0 ± 0.71% of ZsGreen^+^ cells, n = 3 mice) and approximately 70% of Axin2^+^ cells did not express *Ki67* (Fig. 1g, *Ki67*^+^; 26.5 ± 4.4% of ZsGreen^+^ cells, n =3), suggesting that Axin2^+^ cells represent slow-cycling cells. We then examined the expression of *Nt5e* (*Cd73*) and *Pthlh* (*Pthrp*), which are previously-reported stem cell markers of resting chondrocytes^4,5^ A large population of Axin2^+^ cells expressed *Cd73* (Fig. 1h, *Cd73*^+^;75.8 ± 14.2% of ZsGreen^+^ cells, n=3 mice), and approximately one-third of Axin2^+^ cells expressed *Pthrp* (Fig. 1i, *Pthrp*^+^;33.3 ± 7.34% of ZsGreen^+^ cells, n=3 mice). *Foxa2*-expressing cells have been identified to be present close to SOC in the resting zone and to contribute to growth plate renewal.^19^ At P26, *Foxa2* transcripts were sparsely expressed in the resting and proliferative chondrocytes, but were abundantly expressed in the hypertrophic chondrocytes (Fig. S3a). The immunoreactive cells to the anti-Foxa2 antibody were detected at the border of SOC, with some of the cell being ZsGreen positive (Fig. S3b, arrowhead). These data suggest that the identified Axin2^+^ cells included a subset of previously reported chondroprogenitors.

### Self-renewal and Differentiation of Axin2^+^ Cells

To determine the self-renewal ability of the Axin2^+^ cells and their contribution to growth plate maintenance, a series of cell tracing analyses were performed in the *Axin2Cre^ERT2^;R26R^ZsGreen^* mice. After remaining in the resting zone (P26, Fig. 2a), the descendants of initially labeled Axin2^+^ cells (hereafter, *Axin2-creER^+^* cells) first formed short columns (composed of < 10 cells) after 7 days of chase (P30, Fig. 2b, arrowhead) and subsequently formed long columns (composed of ≥ 10 cells) after 10 days of chase (P33, Fig. 2c, arrow). After two weeks of chase, *Axin2-creER+* cells had constituted the columns encompassing from the resting to hypertrophic zones (P37, Fig. 2d, proximal tibia and Fig. S1, distal femur). The number of ZsGreen+ cells in the top 50 μm zone increased during the first month of chase and decreased thereafter (Fig. 2a–e, g). The number of short ZsGreen^+^ columns increased from P23 to P30 and then plateaued, whereas long ZsGreen^+^ columns appeared at P33 and their number increased over time (Fig. 2h). *Axin2-creER^+^* cells continued to accumulate over a long chase period (Fig. 2i) and formed chondrocyte columns in the growth plate at least after six months of tracing (Fig. 2f). We analyzed the data of lineage tracing experiments in male and female *Axin2Cre^ERT2^;R26R^ZsGreen^* mice because there were no significant sex differences in the number of ZsGreen+ cells in the top 50 μm zone, the number of short/long ZsGreen^+^ columns, and the percentage of ZsGreen^+^ cells among the growth plate chondrocytes (Fig. S4a–d).

**Figure 2.**
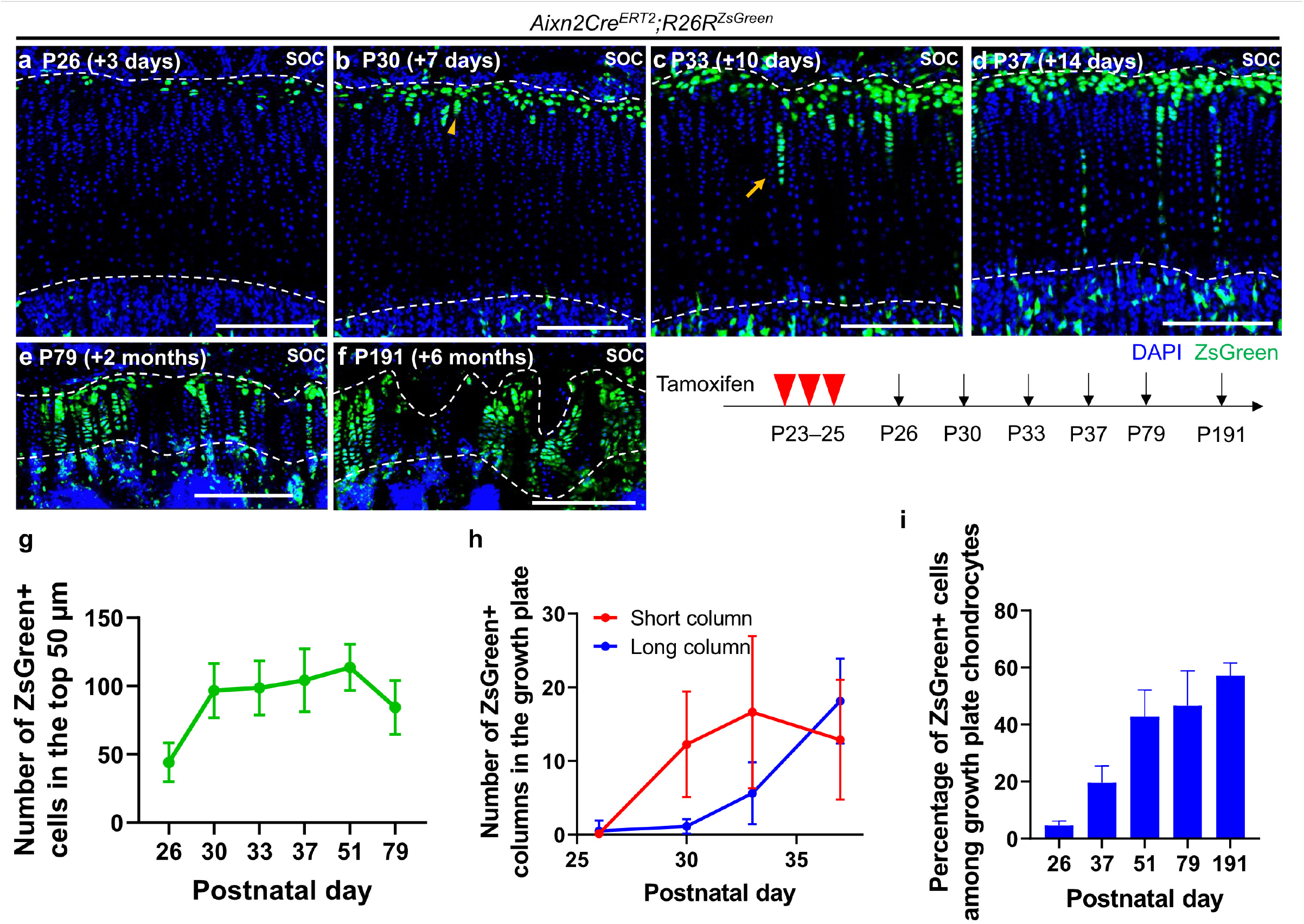
Axin2^+^ cells include a subset of chondroprogenitors in the growth plate. **a–f** Fate-mapping analysis of Axin2^+^ cells in the proximal tibial growth plate in *Axin2Cre^ERT2^;R26R^ZsGreen^* mice (pulsed on P23–25, and traced for several days). Representative images showing descendants of initially labeled Axin2^+^ cells at 3 (n=8) (**a**), 7 (n=8) (**b**), 10 (n=8) (**c**) and 14 (n=8) (**d**) days, and at 2 (n=7) (**e**) and 6 (n=6) (**f**) months after tamoxifen injection. Arrowhead, short column (≤ 9 cells) (**b**); arrows, long column (≥ 10 cells) (**c**). **g–i** Quantification of ZsGreen+ cells in the top 50 μm (green line) (**g**), ZsGreen+ columns in the growth plate, short columns (≤ 9 cells, red line) and long columns (≥ 10 cells, blue line) (**h**), and percentage of ZsGreen^+^ cells among growth plate chondrocytes (blue bars) (**i**). P26–P51 (n = 8), P79 (n = 7), P191 (n = 6). SOC, secondary ossification center. The white dashed lines demarcate the growth plate from the surrounding tissues. Scale bars: 200 μm (**a–f**). All data are presented as the mean ± SD.

To investigate whether ZsGreen^+^ chondrocyte columns existed in the same clone, we analyzed *Axin2Cre^ERT2^;R26R^Confetti^* mice. The administration of tamoxifen for five consecutive days from P23 marked Axin2^+^ cells in the resting zone (Fig. S5a, orange arrow heads), enabling us to perform clonal analyses. After three months of chase, some *Axin2-creER^+^* cells formed chondrocyte columns labeled in the same color (Fig. S5b, orange arrow and Fig. S5d, magenta, 23.3 ± 6.35%), indicating that single Axin2^+^ resting chondrocyte can give rise to the entire chondrocyte column. In contrast, some *Axin2-creER^+^* cells remained in singlet (Fig. S5c, white arrow and Fig. S5d, black, 38.9% ± 6.89) or doublet form (Fig. S5c, white arrowhead and Fig. S5d, blue, 34.8 ± 7.14%) in the resting zone, indicating that some Axin2^+^ cells can be dormant or very slow-proliferating resting chondrocytes. Together, these findings showed that Axin2^+^ cells included a subset of chondroprogenitors with self-renewal capability to continually form growth plate chondrocytes during the bone growth period.

We next examined whether Axin2^+^ cells at different developmental stages had the capacity to form chondrocyte columns. The Axin2^+^ cells at both P45 and P87 were found in the resting zone (Fig. 1b–d and Fig. S6a–c) in a similar distribution to that observed at P26 (Fig. 1b and Fig. S6a). Interestingly, Axin2^+^ cells were also observed in the hypertrophic zone at P87 (Fig. S6c), suggesting that Wnt/β-catenin signaling was highly activated in the hypertrophic zone at this time. After one month of chase analysis, Axin2^+^ cells initially labeled at both P45 and P87 had produced long chondrocyte columns (Fig. S6e, arrows, and Fig. S6f, arrow). The percentage of *Axin2-CreER*^+^ cells among growth plate chondrocytes after 28 days of chase decreased with time (Fig. S6d–g). These data suggested that Axin2+ cells in the resting zone retained the ability to generate chondrocyte columns over time, although their contribution to growth plate chondrocytes declined with age.

### Establishment of Mouse Catch-up Growth Model

To investigate changes in the distributions of *Axin2-CreER*^+^ cells during malnutrition and subsequent catch-up growth, we first developed a mouse catch-up growth model. The mice were fed ad libitum from the start (control group), or subjected to a 50% dietary restriction (DR) for seven days from P27 before ad libitum feeding (catch-up group) (Fig. S7a). During the seven-day DR period, male and female catch-up group mice gained less weight than control group mice (male, p = 0.0023, and female, p = 0.0120; Fig. S7b, c). During the first five days after DR, male and female mice in the catch-up group gained weight more rapidly than control mice (male, p = 0.0005, and female, p = 0.0259; Fig. S7b, c), and there were no significant differences in body weight at the end of the experiment for both males and females (Fig. S7b, c).

Tibial length of male mice in the catch-up group was 0.64 mm less than that of control animals at P34 (p = 0.0186; Fig. S7d). Progressive catch-up growth occurred (p = 0.0008), reducing this deficit to 0.02 mm (not significant) at P62 (Fig. S7d). Similarly, tibial length of female mice in the catch-up group was 0.75 mm less than that of control animals at P34 (p = 0.0163; Fig. S7e), and progressive catch-up growth occurred (p = 0.015), reducing this deficit to 0.10 mm (not significant) at P62 (Fig. S7e). Proximal tibial growth rates, measured by alizarin labeling, were significantly reduced by seven-day DR compared with those of the control groups for both male and female (p < 0.0001, and p = 0.0039, respectively; Fig. 4f–h). Marked increases were observed in the growth rate of catch-up group mice between P34 and P41 in both males and females (p = 0.0095, and p = 0.0166, respectively; Fig. S7f–g). The growth rate in the catch-up group then gradually decreased but remained greater than that of the control group, at least until P48, in both males and females (Fig. S7f–g). The *Axin2Cre^ERT2^;R26R^ZsGreen^* mice showed similar reduction in the growth rate during seven-day DR compared to the control group (p<0.0001), and increasein the growth rate after refeeding (p=0.0119) (Fig. S8a).

Histomorphometry analysis revealed that the DR reduced the lengths of total growth plate and hypertrophic zone (Fig. S7h). However, the DR increased the ratio of the length of the resting zone to that of the total growth plate, whereas the DR did not change the ratio of the length of other zones to that of the total growth plate (Fig. S7h). During the catch-up growth, the growth plate length was significantly increased compared to the ad libitum control after stopping DR (Fig. S7i). The ratio of the hypertrophic zone to the total growth plate length was not significantly different between these two groups at any time points (Fig. S7i). The EdU incorporation rate in the growth plate was markedly reduced by DR and was recovered in 2 days after stopping DR(Fig. S7j).

### DR Enhances Self-replication of Resting Chondrocytes While Inhibiting Their Differentiation

The behavior of chondroprogenitors during DR and following catch-up growth was investigated by applying the lineage tracing analysis in *Axin2Cre^ERT2^;R26R^ZsGreen^* mice to the mouse catch-up growth model. First, we examined the effect of seven-day DR on the distribution of *Axin2-CreER*^+^ cells in the proximal tibial growth plate. After tamoxifen administration for three consecutive days from P23, the mice were fed ad libitum (control group) or subjected to seven-day DR (DR group) (Fig. 3a). After 11 days of tracing (P34), *Axin2-CreER*^+^ cells had formed fewer chondrocyte columns in the DR group than in the control counterparts (Fig. 3b–d). In support of this, histological analysis revealed that the growth plates of DR mice showed reduced chondrocyte column density compared with those of the control mice (Fig. 3i–k). DR led to a 30% increase in the number of ZsGreen+ cells in the top 50 μm zone compared to that in the control group (Fig. 3b, c, e). To independently confirm this observation, we performed *in situ* hybridization of *Clusterin* (*Clu*), of which expression has been detected in the resting zone as well as in the articular cartilage,^20^ to quantify the number of resting chondrocytes. We chose *Clu* as a marker for resting chondrocytes because *Clu* expression was more abundant and more selective for resting chondrocytes compared to the previously-reported markers for resting chondrocytes, including *Cd73*, *Pthlp* and *Foxa2*(Fig. S9a, b). The number of *Clu*^+^ resting chondrocytes was 56% higher in DR mice than in control mice (Fig. 3f–h).

**Figure 3.**
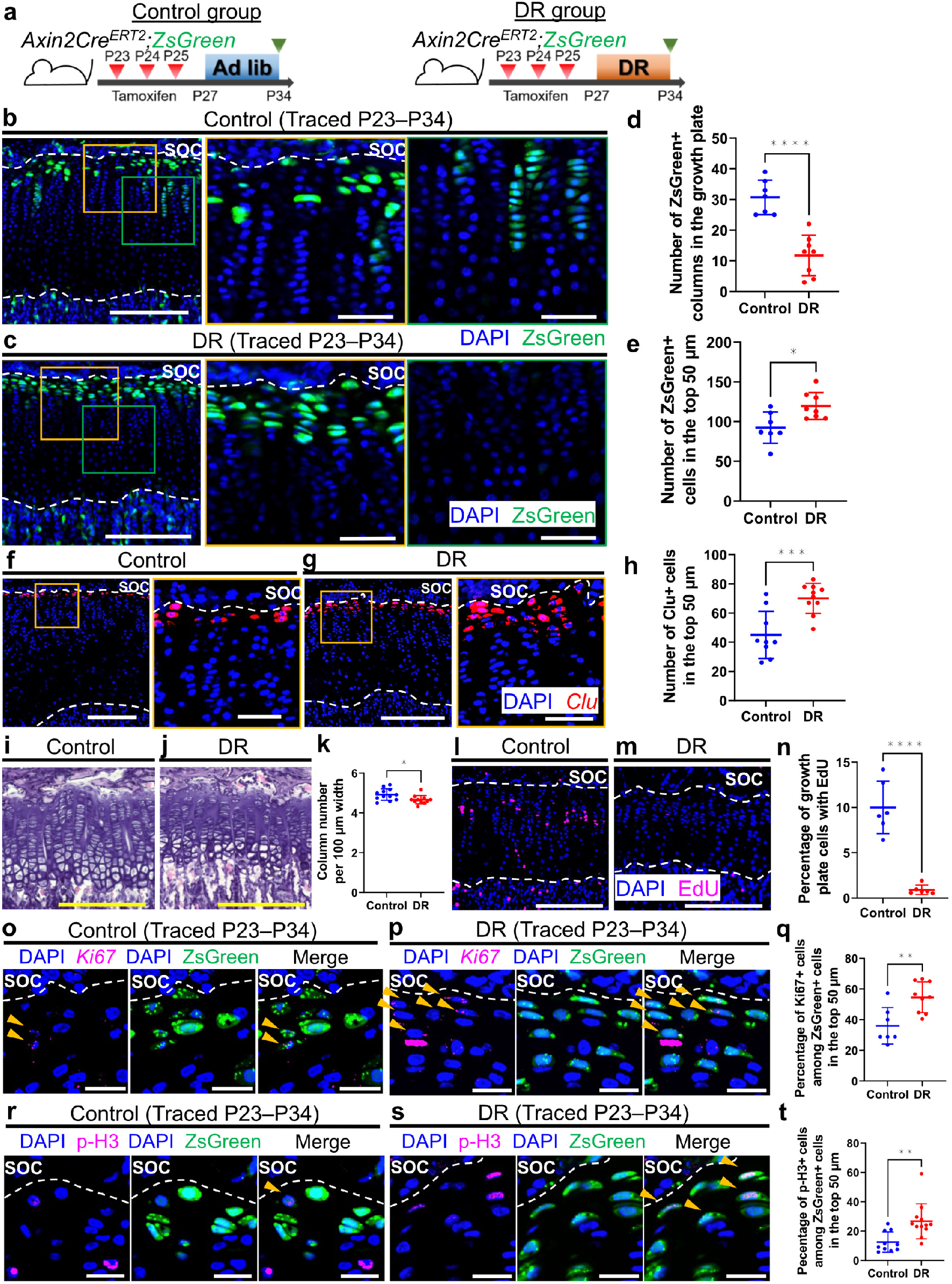
Dietary restriction enhances the self-replication of resting chondrocytes while inhibiting their differentiation. **a** Schematic diagrams of the fate-mapping analysis of Axin2^+^ cells in the proximal tibial growth plate in *Axin2Cre^ERT2^;R26R^ZsGreen^* mice (pulsed on P23–25, and traced for 11 days). The mice were fed ad libitum (control group) or were subjected to seven-day DR from P27 (DR). **b**, **c** Representative images of descendants of initially labeled Axin2^+^ cells (ZsGreen^+^ cells) in the proximal tibial growth plate in the control (**b**) and DR (**c**) groups. Middle panels show magnified views of resting zone of the growth plate, and right panels show magnified views of proliferative zone of the growth plate. **d**, **e** Quantification of the ZsGreen^+^ columns in the growth plate (**d**) and ZsGreen^+^ cells in the top 50μm (**e**). Control group (n = 7), DR group (n = 9). ****p = < 0.0001 (**d**), *p = 0.0125 (**e**). **f**, **g** Representative images for *in situ* hybridization of *Clu* in the proximal tibial growth plate in the control (**f**) and the DR (**g**) groups. Right panels show magnified views of the resting zone of the growth plate. **h** Quantification of *Clu* + cells in the top 50 μm. Control group (n = 9), DR group (n = 10). ***p = 0.0008. **i**, **j** Representative images of hematoxylin and eosin staining of the proximal tibial growth plate in the control (**i**) and the DR (**j**) groups. **k** Quantification of column number per 100 μm width of the growth plate. Control group (n = 12), DR group (n = 11). *p = 0.0245. **l**, **m** Representative images of EdU labeling in the proximal tibial growth plate in the control (**l**) and the DR (**m**) groups. EdU (50 μg/g body weight) was injected two hours before euthanization. **n** Percentage of growth plate cells with EdU. Control group (n = 6), DR group (n = 6). ****p = < 0.0001. **o**, **p** Representative images for *in situ* hybridization of *Ki67* and ZsGreen^+^ cells in the proximal tibial growth plate in the control (**o**) and the DR (**p**) groups. Arrow heads, ZsGreen^+^ cells expressing *Ki67* (**o**, **p**). **q** Percentage of *Ki67*^+^ cells among ZsGreen^+^ cells in the top 50 μm. Control group (n = 7), DR group (n = 9). **p= 0.0045. **r**, **s** Representative images for immunostaining for p-H3 and ZsGreen^+^ cells in the proximal tibial growth plate in the control (**r**) and the DR (**s**) groups. Arrow heads, ZsGreen^+^ cells expressing p-H3 (**r**, **s**). **t** Percentage of p-H3 positive cells among ZsGreen+ cells in the top 50 μm. Control group (n = 10), DR group (n = 12). **p= 0.0035. Ad lib, ad libitum, DR, dietary restriction, SOC, secondary ossification center, p-H3, phospho histone H3. The white dashed lines demarcate the growth plate from the surrounding tissues. Scale bars: 200 μm (**b** [left-most], **c** [left-most], **f** [left], **g** [left], **i**, **j**, **l**, **m**), 50 μm (**b** [right two], **c** [right two], **f** [right], **g** [right]), 20 μm (**o**, **p, r, s**). All data are presented as the mean ± SD. Statistical significance was determined by unpaired two-tailed *t*-test.

The observation of an increased number of resting chondrocytes with inhibited differentiation into proliferative chondrocytes indicated that DR enhanced the proliferation of resting chondrocytes while reducing the proliferation of more differentiated proliferative chondrocytes. To test this observation, we examined the incorporation of EdU into the growth plate to assess the proliferation of proliferative chondrocytes. When EdU was administered 2 h before euthanization, DR mice exhibited fewer EdU^+^ cells in the growth plate compared to control mice (10.0% ± 2.9% versus 0.9% ± 0.5%; Fig. 3l–n). Because there was little EdU uptake into resting chondrocytes, we performed *in situ* hybridization of *Ki67* and immunostaining for phospho histone H3 (p-H3) to assess the proliferation of resting chondrocytes. The results showed that DR mice showed a higher percentage of *Ki67*^+^ cells among ZsGreen^+^ resting chondrocytes (40.8% ± 11.1% versus 55.8% ± 9.9%; Fig. 3o–q) and higher percentage of p-H3 positive cells among ZsGreen+ resting chondrocytes (12.6% ± 6.9% versus 26.8% ± 12.0%; Fig. 3r–t), suggesting that the resting chondrocytes may increase mitotic activity during DR. Both the decrease in the number of resting chondrocytes exiting to the proliferative zone and the increase in the number of resting chondrocytes suggest that DR alters the balance between the self-expansion and differentiation of chondroprogenitor cells.

### Ad Libitum Feeding After DR Promotes Committed Differentiation of Resting Chondrocytes

Next, we assessed the changes in the distribution of *Axin2-CreER*^+^ cells during the catch-up growth phase after DR. We compared the number of ZsGreen+ cells in the top 50 μm and the number of ZsGreen^+^ columns in mice which were fed ad libitum after 7-day DR (catch-up group) with those in mice under continued-DR (DR group) (Fig. 4a). Two days after ending DR (P36), *Axin2-CreER*^+^ cells constituted a greater number of chondrocyte columns in the catch-up group than in the DR group (Fig. 4b, c, f), suggesting that the resting chondrocytes restarted differentiation into proliferative chondrocytes within 2 days after DR cessation. There were no significant differences in the number of ZsGreen+ chondrocytes in the top 50 μm between the groups at this time point (Fig. 4b, c, g). Seven days after DR cessation (P41), *Axin2-CreER*^+^ cells in the catch-up group had formed a greater number of ZsGreen^+^ columns compared to those formed at 2 days after DR cessation (P36) (Fig. 4c, e, f), whereas the number of ZsGreen^+^ columns remained unchanged in the DR group (Fig. 4b, d, f). This suggested that the pooled resting chondrocytes caused by seven-day DR retained the capacity to make chondrocyte columns. Seven-day feeding after DR cessation led to a 17% decrease in the number of ZsGreen^+^ cells in the top 50 μm compared with that in the DR group (Fig. 4d, e, g). These data indicated that ad libitum feeding after stopping DR promoted the committed differentiation of resting chondrocytes.

**Figure 4.**
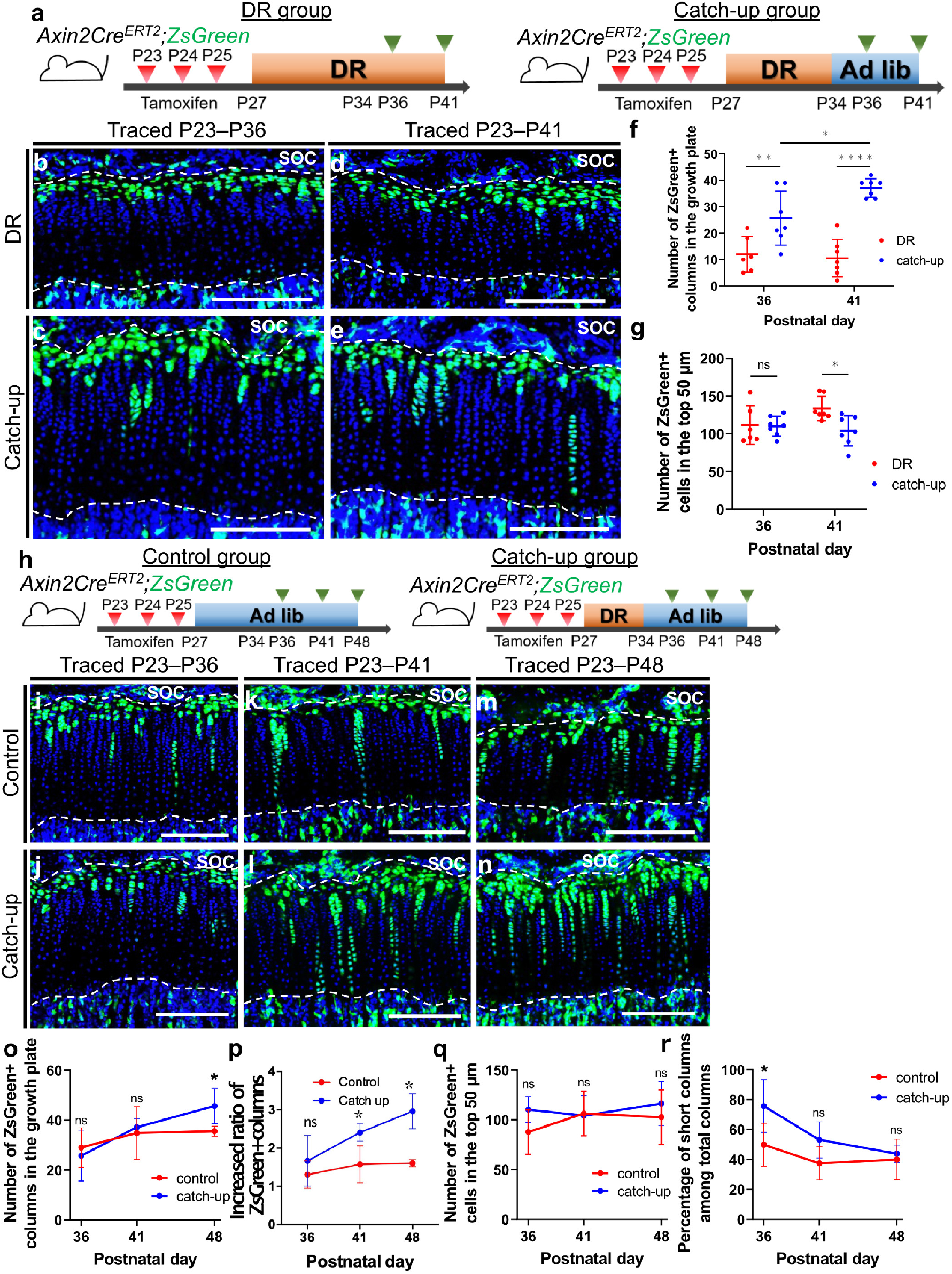
Ad libitum feeding after dietary restriction promotes committed differentiation of resting chondrocytes. **a** Schematic diagrams of the fate-mapping analysis of Axin2^+^ cells in the proximal tibial growth plate in *Axin2Cre^ERT2^;R26R^ZsGreen^* mice (pulsed on P23–25, and traced for 13 or 18 days). The mice were subjected to continued-DR (DR group) or were fed ad lib after seven-day DR (catch-up group). **b–e** Representative images of descendants of initially labeled Axin2+ cells (ZsGreen^+^ cells) in the proximal tibial growth plate in the DR (**b**, **d**) and the catch-up (**c**, **e**) groups. **f**, **g** Quantification of the ZsGreen^+^ columns in the growth plate (**f**) and ZsGreen^+^ cells in the top 50μm (**g**). DR group (n = 6), catch-up group (n = 7) at P36. DR group (n = 7), catch-up group (n = 7) at P41. Adjusted *p = 0.022 (P36 vs P41 in the catch-up group), adjusted **p = 0.0052 (DR vs catch-up, P36), adjusted ****p= < 0.0001 (DR vs catch-up, P41) (**f**), and adjusted p > 0.9999 (ns) (DR vs catch-up, P36), adjusted *p = 0.0157 (DR vs catch-up, P41) (**g**). **h** Schematic of the fate-mapping analysis of Axin2+ cells in the proximal tibial growth plate in *Axin2Cre^ERT2^;R26R^ZsGreen^* mice (pulsed on P23–25, and traced for 13, 18, or 25 days). The mice were fed ad lib (control group) or were fed ad lib after seven-day DR (catch-up group). **i–n** Representative image of ZsGreen+ cells in the proximal tibial growth plate in the control (**i**, **k**, **m**) and the catch-up (**j**, **l**, **n**) groups. **o–r** Quantification of the number of ZsGreen^+^ columns in the growth plate (**o**), the increase rate of ZsGreen+ column from P34 (**p**) and ZsGreen+ cells in the top 50μm (**q**), and percentage of short columns among total columns (**r**). Control group (n =7), catch-up group (n = 7) at P36, P41. Control group (n = 6), catch-up group (n = 6) at P48. Adjusted p > 0.9999 (ns) (P36), adjusted p > 0.9999 (ns) (P41), adjusted *p = 0.0461 (P48) (**o**). Adjusted p=0.3743 (ns) (P36), adjusted **p=0.0043 (P41), adjusted ****p=<0.0001 (**p**). Adjusted p = 0.1865 (ns) (P36), adjusted p > 0.9999 (ns) (P41, P48) (**q**). Adjusted *p = 0.0429 (P36), adjusted p = 0.1003 (ns) (P41), adjusted p > 0.9999 (ns) (P48) (**r**). Ad lib, ad libitum, DR, dietary restriction, ns, not significant, SOC, secondary ossification center. The white dashed lines demarcate the growth plate from the surrounding tissues. Scale bars: 200 μm (**b–e** and **i–n**). All data are presented as the mean ± SD. Statistical significance was determined by two-way analysis of variance and Bonferroni’s multiple comparison test.

The distribution of *Axin2-CreER*^+^ cells in the catch-up group was then compared with that in the ad libitum control group (Fig. 4h). *Axin2-CreER*^+^ cells constituted a significantly higher number of ZsGreen^+^ chondrocyte columns in the catch-up group 14 days after stopping DR (P48) than that in the control group (Fig. 4m, n, o). The DR inhibited differentiation of the resting chondrocytes into proliferative chondrocytes as described above. To assess the response of the resting chondrocytes to refeeding, we examined how much ZsGreen+ columns were increased from P34, completion of the diet restriction, to P36-P48, 2-14 days after refeeding. The average of the increased ratio was significantly higher in the catch-up group at P41 and P48 than the control group (Fig.4p). The increased column formation observed in the catch-up group theoretically contributed to the accelerated growth observed at P41 and P48 in the catch-up growth model (Fig. S7f–h). There were no significant differences in the number of ZsGreen+ cells in the top 50 μm between the groups at all time points examined, although this number tended to increase beginning two days after DR cessation (P36) (p = 0.18) (Fig. 4i–n, q). The percentage of short columns among total columns in the catch-up group was 25.8% higher than that in the control group (75.7 ± 16.2% versus 49.9 ± 13.2%, p = 0.043; Fig. 4i, j, r) two days after stopping DR (P36). Progressive composition changes from short to long columns occurred in the catch-up group during ad libitum feeding (p = 0.0126), reducing this increase to 3.8% (not significant) at 14 days after stopping DR (P48) (Fig. 4r). This indicated that the pool of resting chondrocytes formed during DR comprised a heterogeneous cell population that included both short- and long-term chondroprogenitors. Together, these data demonstrated that external nutrition played an important role in determining the fate of chondroprogenitor cells in the resting zone.

### IGF1/PI3K is Activated in the Resting Zone of the Growth Plate

To understand the molecular mechanism by which external nutritional status regulates the fate of chondroprogenitors, we first investigated signaling pathways that were specifically activated in the resting or proliferative chondrocytes by performing transcriptome analysis using laser microdissection (LMD) and RNA-sequencing (RNA-seq) (Fig. 5a). We microdissected resting chondrocytes labeled with red fluorochrome-labeled and non-labeled proliferative chondrocytes in P30 tibial and femoral growth plates (Fig. S10a–c). Unsupervised clustering analysis showed that resting and proliferative chondrocytes clustered independently (Fig. S10d), indicating that resting chondrocytes had a biologically unique pattern of transcriptomes from those of proliferative chondrocytes. Statistical analysis revealed that 1442 genes were differentially expressed between the two groups (fold change > ± 2, FDR < 0.01), of which 899 and 543 genes were upregulated in resting chondrocytes and proliferative chondrocytes, respectively (Fig. S10e). Representative genes upregulated in resting chondrocytes included previously reported markers for resting chondrocytes, such as *Clu*,^20^ *Gas1*, *Wif1,*^21^ *Efemp1*, and *Sorl1*^22^ (Fig. S10e). Representative genes downregulated in the resting chondrocytes included *Acan, Col2a1, Matn1 and Slc2a1* (Fig. S10e). Downregulation of these genes in the resting chondrocytes was confirmed by *in situ* hybridization(Fig. S10f). The expression of the markers for prehypertrophic chondrocytes, such as *Ihh*,^23^ *Mef2c*,^24^ and *Pth1r*^25^, were uup-regulated in the proliferative chondrocytes (Fig. S10e), which is likely due to the contamination of the prehypertrophic chondrocytes during excision of the proliferative chondrocytes.

**Figure 5.**
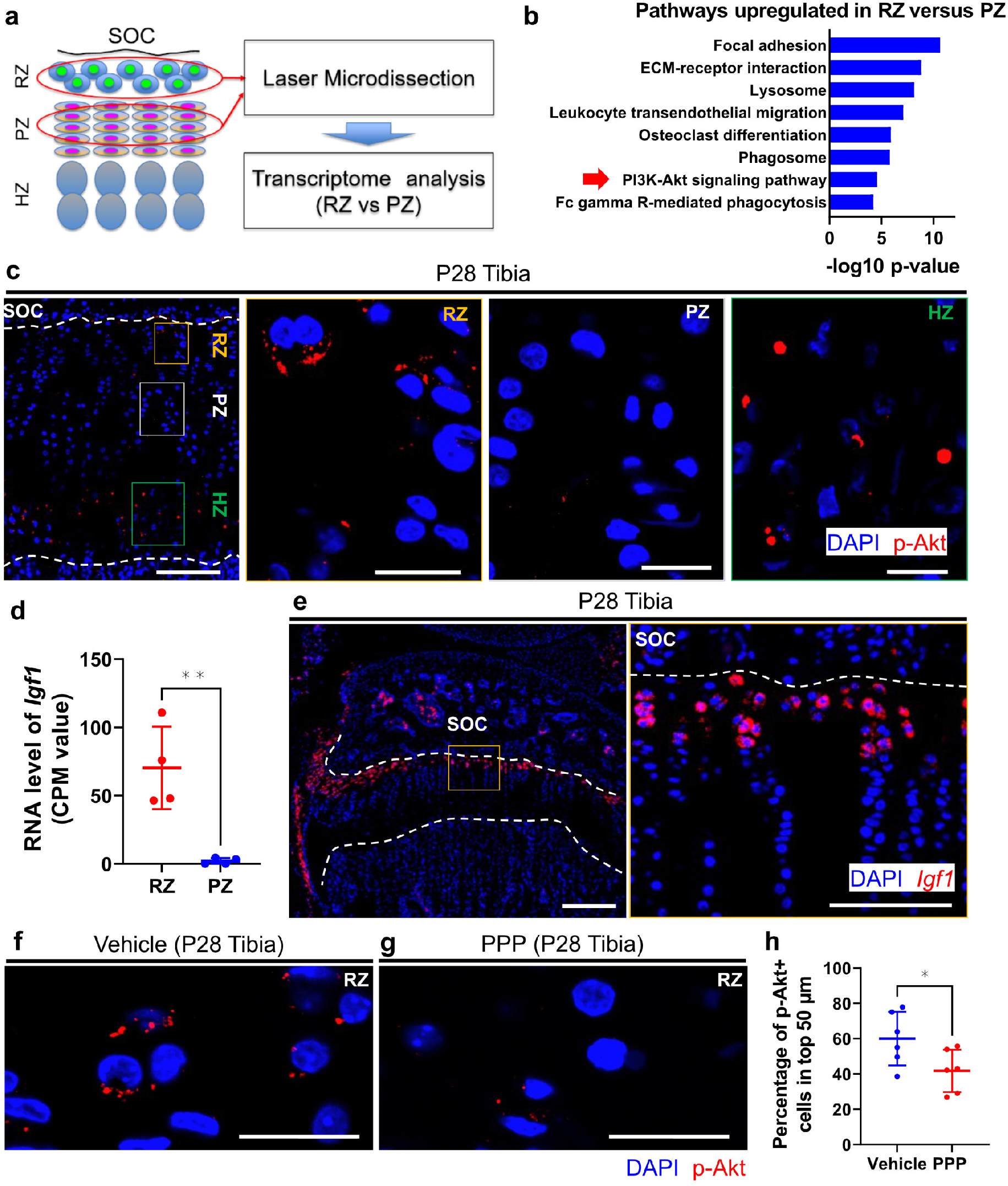
IGF-1/PI3K signaling is activated in the resting zone. **a** Schematic diagram for transcriptome analysis using laser microdissection and RNA sequencing. P30 tibial and femoral growth plates in *Axin2Cre^ERT2^;R26R^TdTomato^* mice (pulsed on P23–25) were subjected to laser microdissection. **b** KEGG pathways enriched by genes significantly upregulated in the resting zone compared to the proliferative zone. Red arrow indicates PI3-Akt signaling pathway. Logarithmic p-values of significance are indicated on the *x*-axis. **c** Representative image of immunohistochemistry for p-Akt in the proximal tibial growth plate at P28. Right three panels show magnified views of each zone of the growth plate. **d** CPM values of *Igf-1* of the resting and proliferative zone samples. RZ (n = 4), PZ (n = 4). **p = 0.0041. **e** Representative image for *in situ* hybridization of *Igf-1* in the proximal tibial growth plate at P28. Right panel shows the magnified view of resting zone of the growth plate. **f**, **g** Representative image of immunohistochemistry for p-Akt in the resting zone of the proximal-tibial growth plate at P28 after receiving either vehicle (dimethyl sulfoxide/ corn oil, 9:1) (**f**) or IGF-1 receptor tyrosine kinase inhibitor (PPP, 20 ug/g body weight) (**g**) 30 minutes before euthanization. **h** Percentage of p-Akt+ cells in the top 50 μm. Vehicle group (n = 6), PPP group (n = 6). *p = 0.0435. p-AKT, phosphorylation of the protein kinase Akt, SOC, secondary ossification, RZ, resting zone, PZ, proliferative zone, HZ, hypertrophic zone, CPM, count per million, PPP, picropodophyllin. The white dashed lines demarcate the growth plate from the surrounding tissues. Scale bars: 200 μm (**c** [left-most], **e** [left]), 100 μm (**e** [right]), 20 μm (**c** [right three], **f**, **g**). All data are presented as the mean ± SD. Statistical significance was determined by unpaired two-tailed *t*-test.

Pathway analysis of differentially expressed genes (DEGs) revealed significant enrichment of several Kyoto Encyclopedia of Genes and Genomes (KEGG) pathways. KEGG pathways related to metabolism were often downregulated in resting chondrocytes, including metabolic pathways (KEGG: 01100), fatty acid metabolism (KEGG:01212), glycolysis /gluconeogenesis (KEGG: 00010), biosynthesis of unsaturated fatty acids (KEGG:01040), and biosysthesis of amino acids (KEGG: 01230) (Fig. S10g). These data indicated that resting chondrocytes showed unique metabolic regulations compared to proliferative chondrocytes. Among the pathways upregulated in resting chondrocytes, we focused on the phosphatidylinositol-3-kinase (PI3K) signaling pathway (KEGG:04151) (Fig. 5b) because it is affected by nutrient availability ^26^ and is required for normal growth plate differentiation and endochondral bone growth.^27^ To confirm whether PI3K signaling was activated in resting chondrocytes, the localization of phosphorylation of the protein kinase Akt (p-Akt), a downstream target of the PI3K pathway,^28^ was examined. Immunohistochemistry of p-Akt showed that it was localized specifically in the resting and hypertrophic zones of the growth plate (Fig. 5c). Examination of upstream regulators through Ingenuity Pathways Analysis revealed more activated insulin-like growth factor 1 (IGF-1) signaling in resting chondrocytes than in proliferative chondrocytes (Table S1). Because the PI3K pathway is known as the major signaling cascade downstream of IGF-1 in many cell types,^29–32^ these bioinformatics data suggested that IGF-1/PI3K signaling was activated in resting chondrocytes. Accordingly, the RNA-seq data showed significantly increased expression of *Igf-1* in the resting chondrocytes compared with that in the proliferative chondrocytes (Fig. 5d). *In situ* hybridization also demonstrated that the *Igf-1* mRNA was localized in the resting zone of the growth plate, as well as in the perichondrium (Fig. 5e), and that *Igf1r* mRNA were ubiquitously expressed in the growth plate chondrocytes including Axin2+ cells in the resting zone (Fig. S11, arrowheads). Administration of picropodophyllin, an IGF-1 receptor tyrosine kinase inhibitor, reduced the percentage of p-Akt^+^ cells in the resting zone (Fig. 5f–h), suggesting that the activated PI3K pathway in the resting chondrocytes was at least partially dependent on IGF-1 signaling.

### IGF-1/PI3K Signaling in Resting Chondrocytes Changes in Sync with Nutritional Availability

Because both IGF-1 and PI3K signaling are affected by nutritional status,^26,33,34^ we investigated possible changes in the activity of IGF-1/PI3K signaling during DR and catch-up growth. We compared the serum levels of IGF-1 and the percentage of *Igf-1*^+^ and p-Akt^+^ cells among ZsGreen^+^ resting chondrocytes in catch-up group, nine-day DR mice (DR group), and mice fed ad libitum (control group) (Fig. 6a). Serum IGF-1 levels were significantly reduced in the DR group compared to those of the control and were recovered to some extent in the catch-up group (Fig. 6b), suggesting that circulating IGF-1 levels changed depending on the nutritional status as reported previously.^35^ *In situ* hybridization revealed that local *Igf-1* expression was significantly reduced in the DR group compared to that of the control, and was recovered to some extent in the catch-up group (Fig. 6c–f). The reduction of Igf1 expression in the DR group and its recovery in the catch-up group were confirmed at the protein level (Fig. S12). Similarly, the activity of PI3K signaling, as assessed by p-Akt expression, was significantly reduced by nine-day DR compared to that of the control group, and recovered to control levels in the catch-up group (Fig. 6g–j). These data indicated that both endocrine and para/autocrine IGF-1 and PI3K signaling activity changed with nutritional status. To examine the causal relationship between reduced IGF-1 and PI3K signaling activity under DR, we tested whether rhIGF-1 administration reversed the reduction of PI3K signaling caused by nutritional deprivation. One day of fasting caused a significant decrease in the activity of PI3K signaling and decreased circulating IGF-1 and growth plate *Igf-1* levels compared to control values (Fig. 6k–p, r). Remarkably, this reduction in PI3K was ameliorated by administration of rhIGF-1 (Fig. 6q, r), suggesting the requirement of IGF-1 reduction for downregulation of PI3K signaling caused by nutrient deprivation.

**Figure 6.**
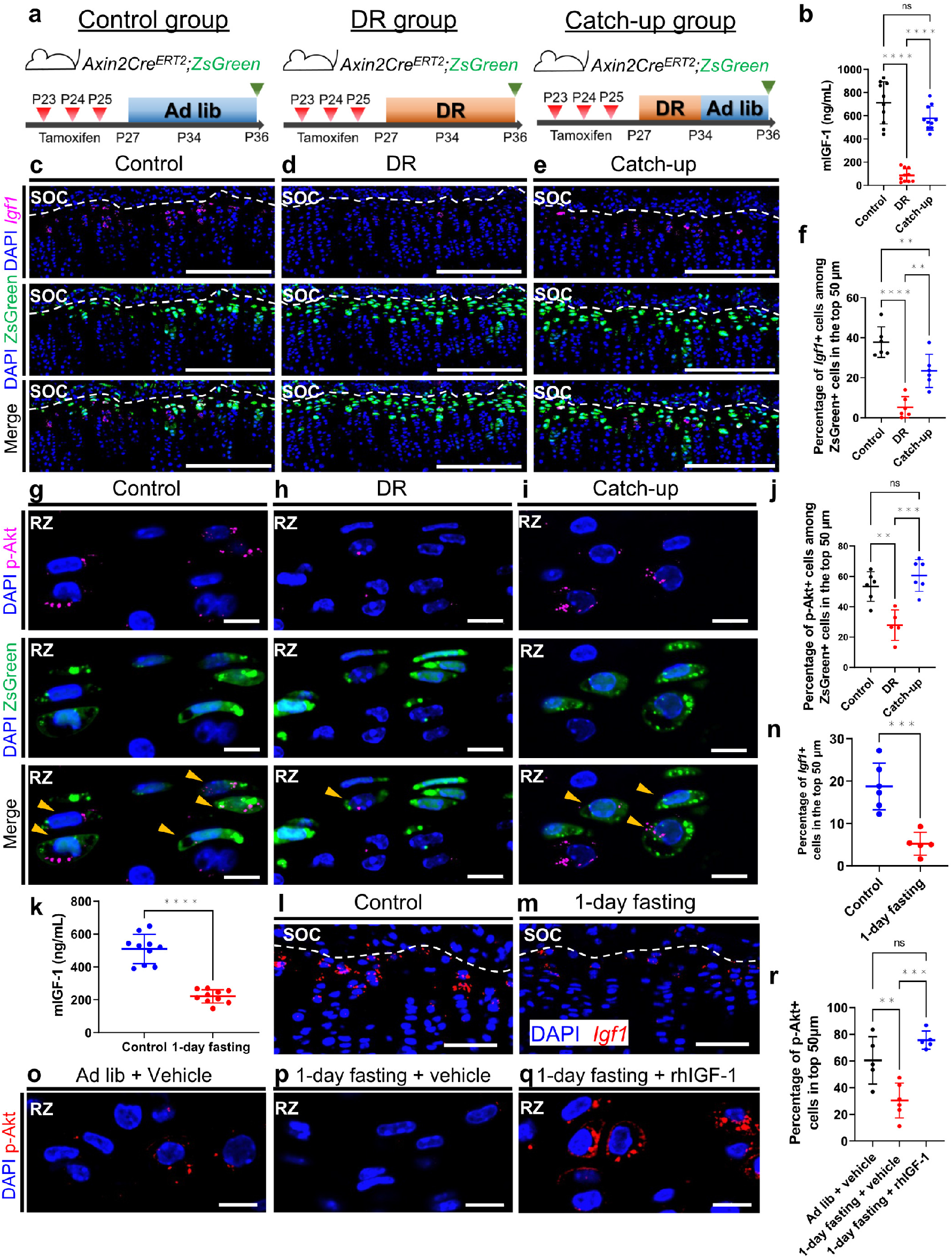
IGF-1/PI3K signaling in resting chondrocytes changes in sync with nutritional availability. **a** Schematic diagrams of the fate-mapping analysis of Axin2^+^ cells in the proximal tibial growth plate in *Axin2Cre^ERT2^;R26R^ZsGreen^* mice (pulsed on P23–25, and traced for 13 days). The mice were fed ad lib (control group), subjected to continued-DR (DR group) or were fed ad lib after 7-day DR (catch-up group). **b** Measurements of circulating IGF-1 levels in the sera from the control (n = 10), DR (n = 10), and catch-up (n = 10) groups. Adjusted ****p < 0.0001 (control vs DR), adjusted ****p < 0.0001 (DR vs catch-up), adjusted p = 0.0631 (ns) (control vs catch-up). **c–e** Representative images for *in situ* hybridization of *Igf-1* and descendants of initially labeled Axin2^+^ cells (ZsGreen^+^ cells) in the proximal tibial growth plate in the control (**c**), DR (**d**), and catch-up (**e**) groups. **f** Percentage of *Igf-1* + cells among ZsGreen+ cells in the top 50 μm. Control group (n = 6), DR group (n = 6), catch-up group (n = 6). Adjusted ****p < 0.0001 (control vs DR), adjusted **p = 0.0015 (DR vs catch-up), adjusted **p = 0.0095 (control vs catch-up). **g–i** Representative images of immunohistochemistry for p-Akt and ZsGreen^+^ cells in the resting zone of the proximal tibial growth plate in the control (**g**), DR (**h**), and catch-up (**i**) groups. **j** Percentage of p-Akt^+^ cells among ZsGreen^+^ cells in the top 50 μm. Control group (n = 6), DR group (n = 6), catch-up group (n = 6). Adjusted **p = 0.0027 (control vs DR), adjusted ***p = 0.0003 (DR vs catch-up), adjusted p = 0.4384 (ns) (control vs catch-up). **k–n** 28-day-old mice were fed ad libitum (control group) or subjected to one-day fasting (1-day fasting group). **k** Measurements of circulating IGF-1 levels in the sera of the control and one-day fasting groups. Control group (n = 10), 1-day fasting group (n = 10). ****p < 0.0001. Data are presented as the mean ± SD. **l**, **m** Representative images for *in situ* hybridization of *Igf-1* in the proximal tibial growth plate in the control group (n = 6) (**l**) and the 1-day fasting group (n = 5) (**m**). **n** Percentage of *Igf-1 +* cells in the top 50 μm. ***p = 0.0008. **o–r** 29-day-old mice received vehicle (PBS) (vehicle group), vehicle followed by 1-day fasting (1-day fasting + vehicle group), or rhIGF-1 (1μg/g body weight) followed by 1-day fasting (1-day fasting + rhIGF-1 group). PBS or rhIGF-1 were administered one hour before euthanization. Representative images of immunohistochemistry for p-Akt in the resting zone of the proximal tibial growth plate at P29 in the vehicle group (n = 5) (**o**), 1-day fasting + vehicle group (n = 6) (**p**) and 1-day fasting + rhIGF-1 group (n = 5) (**q**). Percentage of p-Akt^+^ cells in the top 50 μm (**r**). Adjusted **p = 0.0066 (ad lib + vehicle vs 1-day fasting + vehicle), adjusted ***p = 0.0002 (1-day fasting + vehicle vs 1-day fasting + rhIGF-1), adjusted p = 0.2079 (ns) (ad lib + vehicle vs 1-day fasting + rhIGF-1). Ad lib, ad libitum, DR, dietary restriction, ns, not significant, p-AKT, phosphorylation of the protein kinase Akt, PBS, phosphate-buffered saline, SOC, secondary ossification, RZ, resting zone. The white dashed lines demarcate the growth plate from the surrounding tissues. Scale bars: 200 μm (**c–e**), 50 μm (**l**, **m**), 10 μm (**g–i**, **o–q**). All data are presented as the mean ± SD. Statistical significance was determined by one-way analysis of variance and Tukey’s multiple comparison test (**b**, **f**, **j**, **r**) or by unpaired two-tailed *t*-test (**k**, **n**).

### Exogenous IGF-1 Promotes Committed Differentiation of Resting Chondrocytes Under DR

The finding that IGF-1 signaling in resting chondrocytes changed with nutritional availability indicated that external nutritional cues determined the fate of chondroprogenitors through IGF-1 signaling. To demonstrate that IGF-1 signaling mediates DR-dependent effects on chondroprogenitor cells, we examined whether exogenous IGF-1 could cancel the effect of DR on chondroprogenitor cells. DR mice received rhIGF-1 (1μg/g body weight) or a vehicle as a control for seven consecutive days during DR (Fig. 7a). There were no significant differences in body weight (data not shown), growth plate height, or hypertrophic zone height between two groups (Fig. 7k, l). However, *Axin2-CreER*^+^ cells generated more chondrocyte columns in DR mice treated with rhIGF-1 than in untreated DR mice (Fig. 7b–d), suggesting that inhibited differentiation of resting chondrocytes by DR was partially reversed by IGF-1 treatment. Similarly, histological findings showed that chondrocyte column density increased after rhIGF-1 treatment (Fig. 7i, j, m). Administration of rhIGF-1 led to a 17% reduction in the number of ZsGreen^+^ cells in the top 50 μm compared with that in untreated DR mice (Fig. 7b, c, e). Accordingly, *in situ* hybridization of *Clu* showed that the number of *Clu*^+^ resting chondrocytes was reduced by 26% after administration of rhIGF-1 (Fig. 7f–h). *In situ* hybridization of *Ki67* showed that administration of rhIGF-1 did not affect the percentage of *Ki67*^+^ cells among ZsGreen^+^ resting chondrocytes (Fig. 7n–p). Together, these data demonstrated that the inhibitory effect of DR on the differentiation of resting chondrocytes was at least in part mediated by reduced IGF-1 signaling, and administration of exogenous IGF-1 enhanced the committed differentiation of resting chondrocytes without affecting their proliferation.

**Figure 7.**
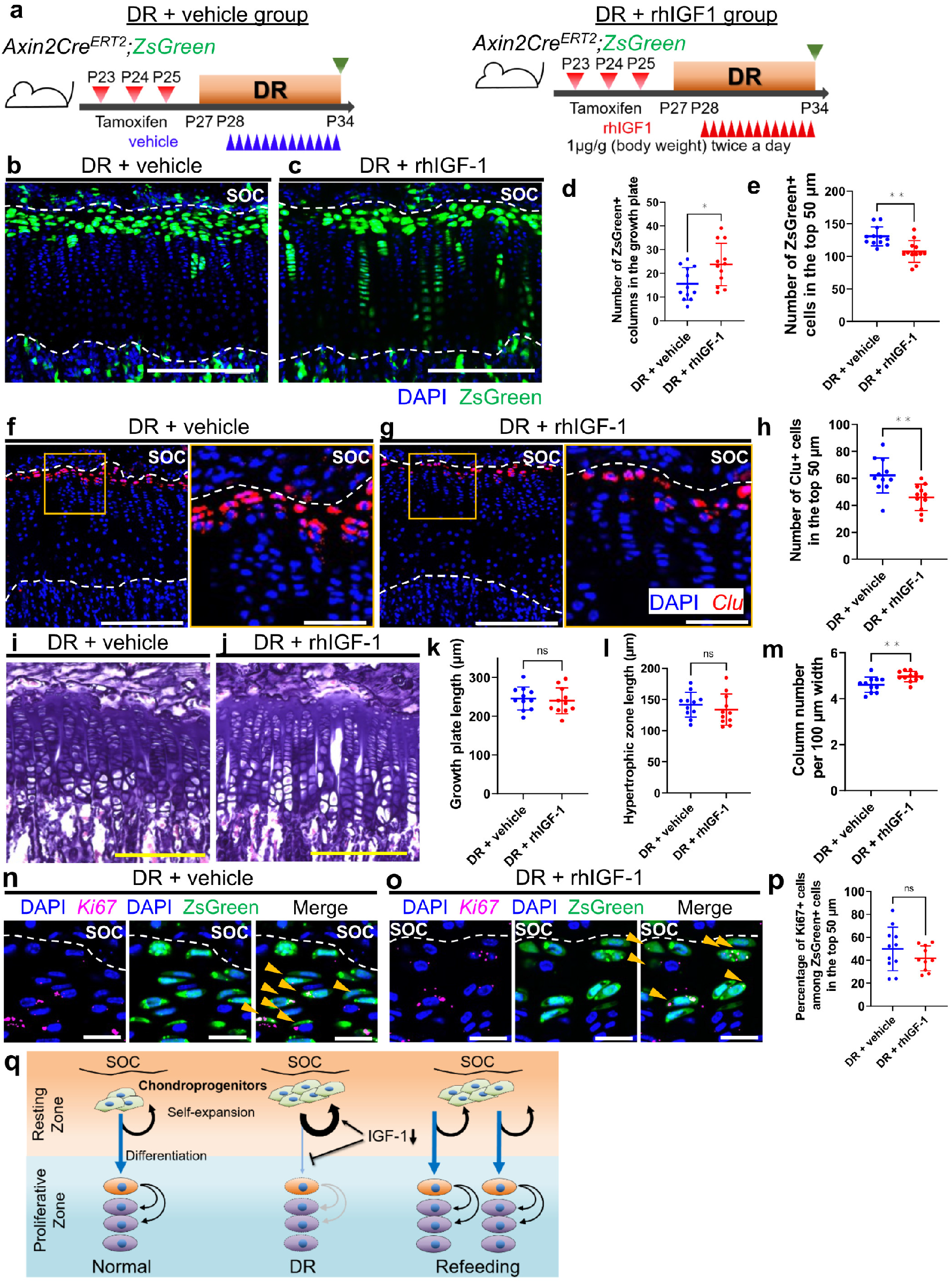
Exogenous IGF-1 promotes committed differentiation of resting chondrocytes under dietary restriction. **a** Schematic diagrams of the fate-mapping analysis of Axin2^+^ cells in the proximal tibial growth plate in *Axin2Cre^ERT2^;R26R^ZsGreen^* mice (pulsed on P23–25, and traced for 11 days). The mice were subjected to seven-day DR with vehicle (PBS) injection (DR + vehicle group) or with rhIGF-1 injection (DR + rhIGF-1 group). Vehicle or rhIGF-1 were administered twice daily for seven days from P28. **b** Representative images of descendants of initially labeled Axin2^+^ cells (ZsGreen^+^ cells) in the proximal tibial growth plate in the DR + vehicle (**b**) and DR + rhIGF-1 (**c**) groups. **d**, **e** Quantification of the ZsGreen+ columns in the growth plate (**d**) and ZsGreen+ cells in the top 50 μm (**e**). DR + vehicle group (n = 12), DR + rhIGF-1 group (n = 12). Statistical significance was determined by unpaired two-tailed *t*-test. *p = 0.0199, **p= 0.0014. Data were presented as the mean ± SD. **f**, **g** Representative images for *in situ* hybridization of *Clu* in the proximal tibial growth plate in the DR + vehicle (**f**), and in the DR + rhIGF-1 (**g**) groups. **h** Quantification of Clu^+^ cells in the top 50 μm. DR + vehicle group (n = 11), DR + rhIGF-1 group (n = 11). Statistical significance was determined by unpaired two-tailed *t*-test. **p = 0.0033. Data are presented as the mean ± SD. **i**, **j** Representative images of hematoxylin and eosin staining in the proximal tibial growth plate in the DR + vehicle (**i**) and the DR + rhIGF-1 (**j**) groups. **k–m** Quantification of growth plate length (**k**), hypertrophic zone length (**l**) and column number per 100 μm width (**m**). DR + vehicle group (n = 11), DR + rhIGF-1 group (n = 11). Statistical significance was determined by unpaired two-tailed *t*-test. p = 0.6669 (ns) (**k**), p = 0.4264 (ns) (**l**), **p = 0.0084 (**m**). Data are presented as the mean ± SD. **n**, **o** Representative images for *in situ* hybridization of *Ki67* and ZsGreen^+^ cells in the proximal tibial growth plate in the DR + vehicle group (**n**) and the DR + rhIGF-1 group (**o**). Arrowheads, ZsGreen^+^ cells expressing *Ki67* (**n**, **o**). **p** Percentage of *Ki67+* cells among ZsGreen^+^ cells in the top 50 μm. DR + vehicle group (n = 11), DR + rhIGF-1 group (n = 10). Statistical significance was determined by unpaired two-tailed *t*-test. p= 0.2485 (ns). **q** Graphical summary how the dynamics of chondroprogenitors change during DR and refeeding. PBS, phosphate-buffered saline, DR, dietary restriction, b.w., body weight, SOC, secondary ossification center, ns, not significant. The white dashed lines demarcate the growth plate from the surrounding tissues. Scale bars: 200 μm (**b**, **c**, **f** [left], **g** [left], **i**, **j**], 50 μm [**f** [right], **g** [right]), 20 μm (**n**, **o**). All data are presented as the mean ± SD. Statistical significance was determined by one-way analysis of variance and Tukey’s multiple comparison test (**b**, **f**, **j**, **r**) or by unpaired two-tailed *t*-test (**k**, **n**).

## Discussion

In this paper, we demonstrated cellular dynamics of chondroprogenitors in response to external nutrition. We identified a specific cell population residing within a niche in the resting zone of the growth plate after SOC formation by analysis of the *Axin2Cre^ERT2^;R26R^ZsGreeni^* mice. The Axin2+ cells included a subset of chondroprogenitors that self-renewed and contributed to growth plate maintenance by differentiating into proliferative chondrocytes. Under DR, these chondroprogenitors are pooled in the resting zone and inhibit their differentiation into proliferative chondrocytes. Once nutritional impairment improved, the pooled chondroprogenitors immediately resumed their differentiation and formed an accelerated rate of chondrocyte columns compared to the control, contributing to catch-up growth. In addition, we showed that nutritional deprivation reduced the activity of IGF-1 signaling in chondroprogenitors, and changed the balance between differentiation and self-expansion of chondroprogenitor cells (Fig. 7q).

Fate-mapping analyses using *Axin2Cre^ERT2^* mice enabled us to directly visualize the dynamics of chondroprogenitors during DR and subsequent catch-up growth. It has been proposed that the proliferation of chondroprogenitor cells in the resting zone is temporarily suppressed during malnutrition, thus preserving proliferative potential. Unused proliferative potential leads to catch-up growth after malnutrition improves.^10,36^ Our results, however, suggest that under DR, chondroprogenitor cells entered the cell cycle without differentiating into proliferative chondrocytes, resulting in their accumulation. Considering that these pooled chondroprogenitors immediately restarted differentiation after nutrition was reestablished, accumulation of this pool of chondroprogenitors may have prepared the growth plate for rapid growth once nutrients became available. In support of this, DR has been shown to stimulate stem cell self-renewal across a wide array of tissue types, seen in intestinal,^37^ hematopoietic,^38^ hair follicle,^39^ and neural^40^ stem/progenitor cell. Our lineage tracing analyses using *Axin2Cre^ERT2^* mice revealed a novel cellular mechanism of catch-up growth, suggesting that *Axin2Cre^ERT2^* mice can be used to monitor the dynamics of chondroprogenitors under several growth inhibitory conditions or local injuries.

The present study showed that nutrient deprivation reduced the activity of both endocrine and para/autocrine IGF-1 signaling, leading to a change in chondroprogenitor behavior during DR. IGF-1 signaling is thought to be a candidate molecular mechanism involved in catch-up growth.^41^ IGF-1 levels are increased in cases of human catch-up growth,^42^ and IGF-1 signaling is necessary for catch-up growth in zebrafish in response to oxygen availability.^43^ In addition, *Igf-1* knockout mice showed a specific growth plate phenotype with an expanded resting zone and a narrowed hypertrophic zone,^44^ supporting our observations in this study (Fig. 3j). Previous studies further support the finding that IGF-1 signaling plays an important role in catch-up growth. The results showing that administration of rhIGF-1 did not rescue either the DR-induced growth plate and hypertrophic shortening (Fig. 7i–l) or the increased induction of chondroprogenitor cell cycle entry (Fig. 7n–p) suggest that IGF-1 signaling is not the only pathway involved in nutrient-induced catch-up growth. Other endocrine, paracrine and autocrine factors may regulate multiple cellular events in the growth plate, such as cell replication of the proliferative zone, hypertrophy and calcification in the hypertrophic zone, and transition from cartilage to bone, in growth arrest and catch-up growth in response to nutritional changes. Further studies are needed to identify the specific signaling molecules that underlie catch-up growth.

Endocrine and para/autocrine IGF-1 signaling is a crucial regulator of growth plate function, as it promotes chondrocyte proliferation and hypertrophy.^45–47^ However, the effects of IGF-1 signaling on resting chondrocytes, including whether they are targeted by IGF-1, remain largely unknown. *In vitro* studies using growth plate chondrocytes have yielded controversial results: some report that IGF-1 selectively acts on differentiated proliferative chondrocytes,^48^ whereas others report that all types of growth plate chondrocytes respond to IGF-1 signaling.^49–51^ In an *in vivo* study on hypophysectomized rats, Ohlson et al.^52^ reported that IGF-1 administration did not affect the number of label-retaining cells (slow-cycling cells) in the resting zone, whereas growth hormone increased it. In contrast, Hunziker et al.^53^ have shown that exogenous IGF-1 application shortens the prolonged cell cycle time induced by hypophysectomy, showing a stimulatory effect of IGF-1 on the mitosis of resting chondrocytes. Our findings suggest that IGF-1 signaling is activated in resting chondrocytes, changing with nutrient availability. Furthermore, exogenous IGF-1 promoted the committed differentiation of resting chondrocytes under restricted dietary conditions, indicating that IGF-1 acts on resting chondrocytes *in vivo*. In the present study, IGF-1 treatment reduced the number of resting chondrocytes (Fig. 7b, c, e) without changing their proliferative activity under DR conditions, as assessed by *Ki67* expression (Fig. 7n–p). DR has been shown to increase the number of resting chondrocytes and allow their entry into the cell cycle, whereas hypophysectomy decreases the proliferative activity of resting chondrocytes.^53^ The discrepancies between our data and previous *in vivo* studies are probably due to the different experimental conditions of IGF-1 administration (i.e., DR versus hypophysectomy). Further investigation is needed to understand the regulatory mechanisms of DR-induced enhancement of chondroprogenitor self-replication.

In summary, we elucidated the dynamics of chondroprogenitors and their regulatory mechanisms in response to nutrient availability. The current identification of these unique characters of chondroprogenitor cells in the growth plate has advanced our knowledge base of bone development and homeostasis. Furthermore, these findings may provide a means to develop non-invasive therapy for human growth disorders.

## Materials and Methods

### Animals

All procedures were conducted according to the National Institutes of Health (NIH) guidelines, the Institutional Animal Care and USE Committee (IACUC) of the University of Maryland, Baltimore, protocol 0120006. *Axin2Cre^ERT2^* (JAX018867), *Rosa26-CAG-loxP-stop-loxP-tdTomato* (Ai9:*R26R-tdTomato*, JAX007909), *Rosa26-CAG-loxP-stop-loxP-ZsGreen* (*Ai6:R26R-ZsG*, JAX007906), *Rosa26-CAG-loxP-stop-loxP-Confetti* (*R26R-Confetti*, JAX013731), and C57BL/6J mice were obtained from the Jackson Laboratory. We crossed *Axin2Cre^ERT2^* mice with *R26R-ZsG*, *R26R-tdTomato* or *R26R-Confetti* to create *Axin2Cre^ERT2^;R26R^ZsGreen^*, *Axin2Cre^ERT2^;R26R^tdTomato^* or *Axin2Cre^ERT2^;R26R^Confetti^* mice, respectively. To achieve Cre recombination, 120 μg/g (body weight) of tamoxifen (T5648; Sigma-Aldrich) dissolved in corn oil (C8267; Sigma-Aldrich) was subcutaneously injected for three consecutive days from P0, P23, P42 or P84 in *Axin2Cre^ERT2^;R26R^ZsGreen^* mice, for three consecutive days from P23 in *Axin2Cre^ERT2^;R26R^tdTomato^* mice, and for five consecutive days from P23 in *Axin2Cre^ERT2^;R26R^Confetti^* mice. All mice were housed in a specific pathogen-free conditions and were allowed free access to food (Laboratory Autoclavable Rodent Diet 5010; LabDiet) and water, except during DR, as described later.

### DR and catch-up growth

C57BL/6J mice were used to establish a mouse catch-up growth model, and *Axin2Cre^ERT2^;R26R^ZsGreen^* mice were used to perform lineage-tracing analyses. After six days of acclimatization in solitary cages, the mice were fed ad libitum or subjected to 50% of their normal food intake from P27 for the indicated periods. The normal intake was based on the intake of the C57BL/6J mice fed ad libitum. During DR, food was provided once daily with ad libitum access to water. The mice were fed ad libitum after seven days of DR for catch-up growth.

### Pharmacological modulation *in vivo*

To investigate whether activation of PI3K signaling in resting chondrocytes is dependent on IGF-1 signaling, 28-day-old C57/B6J mice received intraperitoneal injections of either the IGF-1 receptor tyrosine kinase inhibitor picropodophyllin (PPP, 20 μg/g body weight; SLK-S7668, Selleck Chemicals) or vehicle alone (dimethyl sulfoxide [D8418, Sigma-Aldrich]/corn oil, 9:1) 30 minutes before euthanization.^54^ To examine the relationship between IGF-1 and PI3K signaling under nutritional deprivation, 28-day-old C57/B6J mice were fasted for one day, then treated with a subcutaneous injection of either rhIGF-1 (1 μg/g body weight; 291-G1, R&D Systems) or vehicle phosphate-buffered saline (PBS) alone 60 min before euthanization. *Axin2Cre^ERT2^;R26R^ZsGreen^* mice underwent catch-up growth treatment (See “DR and catch-up growth”) and received subcutaneous rhIGF-1 (1 μg/g body weight) or PBS twice daily (9:00 am and 18:00) for seven consecutive days from P28.^55^ Proximal tibias were collected 2 h after the final dose was administered at 9:00 on P34.

### Measurement of tibial length

Radiographic lateral view images of the fixed tibia were taken using a Faxitron X-ray Specimen Radiography System (Hologic) in automatic exposure control mode. Images were analyzed using ImageJ software.

### Histology and imaging

For Hematoxylin and eosin staining, EdU staining, alizarin labeling, immunohistochemistry, and *in situ* hybridization, frozen sections at 5 μm were analyzed using Keyence BZX710 (Keyence) or Nikon CSU-W1 Spinning Disk Confocal Camera (Nikon). For lineage tracing analyses, frozen sections at 150 μm were analyzed using Nikon CSU-W1 Spinning Disk Confocal Camera. Details are provided in supplementary methods.

### LMD and RNA-seq analysis

RNA was extracted from 8-μm snap frozen sections of distal femur and proximal tibial growth plates in *Axin2Cre^ERT2^;R26R^TdTomato^* mice using the Picopure RNA Isolation Kit (KIT0204, Thermo Fisher) as previously described.^56^ Microdissections of resting chondrocytes and proliferative chondrocytes were performed using a Leica LMD 7000 laser microdissection system (Leica Microsystems). RNA was amplified by two rounds of *in vitro* transcription using the Arcturus RiboAmp HS PLUS Kit (KIT0521, Thermo Fisher) and complimentary DNA libraries were prepared using the KAPA RNA HyperPrep Kit (Roche) according to manufacturer instructions. cDNA libraries were sequenced using an Illumina NovaSeq 6000 system. Raw and processed data are available in the Gene Expression Omnibus (GEO) database under the accession number GSE192840. Details are provided in supplemental methods.

### Statistical Analysis

Statistical analyses were performed using Prism GraphPad (version 6/9). All numerical results were presented as mean ± SD from a minimum of three different experiments (the exact number is indicated in the figure legends). For single comparisons, data were tested for statistical significance using an unpaired two-tailed *t*-test. For multiple comparisons, one-way analysis of variance (ANOVA) followed by Tukey’s post hoc test was used. Comparisons between two groups and time points were performed using two-way ANOVA followed by Bonferroni’s multiple comparisons. The *P* values of statistical significance are shown in the respective figures.

## Acknowledgements

We thank L. Veronique (Children’s Hospital of Philadelphia) and Q. Lin (University of Pennsylvania) for invaluable advices. We also thank K. Storm for professional editing of the manuscript. Research reported in this article was partially supported by the National Institute of Arthritis and Musculoskeletal and Skin Diseases of the National Institutes of Health under Award Number R01AR062908 (to MEI), R01AR056837 (to MI), R01AR073181 (to SO), and R21AR077654 (to SO), and Departmental fund of University of Maryland (to MI, ME-I).

## Contributions

TO and MEI devised the project, the main conceptual ideas and proof outline. TO performed a large part of the experiments. JK, YU and MEI performed some experiments. KW and HT helped with the experiments. SO supported the microdissection and animal experiments. TO, JK and MEI analyzed the data, and YU, YO, TS, ST, MI and SO verified the data analysis. YI performed bioinformatics analysis. TO and MEI wrote the manuscript.

