## Supplementary information for "Nutrient-regulated dynamics of chondroprogenitors in the postnatal murine growth plate"

**Supplementary Methods**

**Tissue processing.** For immunohistochemistry or lineage tracing, knees were fixed in freshly prepared 4% paraformaldehyde (PFA) (158127, Sigma-Aldrich) in PBS for 24h at 4 °C, then decalcified in 10% ethylenediaminetetraacetic acid (EDTA) in PBS (97063-730, VWR International) (pH 7.4) for four days at 4 °C. The samples were transferred to 30% sucrose solution (S0389, Sigma-Aldrich) in PBS for 24 h at 4 °C and embedded in OCT compound (4585, Thermo Fisher Scientific). Five or 150 µm sagittal knee sections at the level of the medial plateau were made using a cryostat (CM1950, Leica Biosystems). Five micrometer sections were prepared on an adhesive film (Cryofilm type IIIc, Section-Lab Co. Ltd.) using the Kawamoto film method.[1] For hematoxylin and eosin (H&E) staining, 5-ethynyl-2′-deoxyuridine (EdU) staining, *in situ* hybridization or alizarin labeling, the knee samples were placed into freshly prepared 2% formaldehyde (28906, Thermo Fisher) in sterile PBS for 24 h at room temperature and for three days at 4 °C, then were transferred to a 30% sucrose/0.5% formaldehyde solution in sterile PBS for 24 h at 4 °C. After embedding the samples in OCT compound, 5 µm sections were prepared using Kawamoto film method. For LMD, the samples were directly embedded in OCT compounds, then snap frozen in hexane dry-ice. One or two sections (as determined from the position of medial plateau and posterior cruciate ligament) were used for H&E staining, EdU staining, alizarin labeling, immunohistochemistry, and *in situ* hybridization.

**Confocal microscopy and image analysis.** Detection and confocal imaging for immunohistochemistry, *in situ* hybridization, and lineage tracing, was performed with a Nikon CSU-W1 Spinning Disk Confocal Camera (Nikon), Imaris image analysis software (Oxford Instruments), and ImageJ software. For lineage tracing analyses, the full thickness 150 µm sections of frozen samples were stained with DAPI for 30 min at room temperature. Samples were then mounted in 75% 2,2'-thiodiethanol (166782, Sigma-Aldrich) mounting buffer following a published method.[2] The sections were scanned, and 3D z-stacks were constructed. The images of the center areas of the proximal tibial growth plate, 800 µm in width and 100 μm in depth, were cropped out and analyzed in surpass and slice modes of Imaris. Chondrocyte columns were defined as aggregations comprising of four or more cells. Columns comprising nine or fewer cells were denoted as a short column, and those containing 10 or more cells as long columns. [3] The number of chondrocyte columns was counted manually in 3D using Imaris software. Three serial 2D images with a 10 μm interval per sample were used to quantify the number of ZsGreen$^+$ cells in the top 50μm[4] and the percentage of ZsGreen$^+$ cells among the growth plate chondrocytes. The length of the resting zone changes with age, and has been reported in ranges 40-60 μm at 2-6 weeks of ages.[5] To avoid complexity of defining the resting zone in individual sections, we counted the number of ZsGreen$^+$ cells in the top 50 μm to quantify the ZsGreen$^+$ cells in the resting zone. For the analysis of ZsGreen$^+$ cells in the resting zone at P3, the ZsGreen$^+$ cell rate was calculated in a square area measuring 200 μm per side, in a region where round chondrocytes were observed (Fig.1a). Quantification was performed using ImageJ software. Detection and imaging for H&E staining, alizarin labeling, and EdU labeling were performed using Keyence BZX710 (Keyence) and quantification was performed using a BZ-X Analyzer (Keyence).

**Histological analysis.** Histological analyses were performed using the results of H&E staining, which was performed according to standard protocols. The central area of the proximal tibial growth plate with a width of 600 μm was cropped and analyzed. Growth plate height was measured parallel to the chondrocyte column. The resting and proliferative zones were defined by morphology following the method described previously.[5] The resting zone was defined from the lower margin of the secondary ossification center to the slightly flat doublet chondrocytes before formation of long columns of proliferative chondrocytes. Hypertrophic chondrocytes were defined by a height ≥ 10 μm as reported previously.[5] Growth plate height and hypertrophic zone height were measured at four equally spaced points in the cropped area. Terminal hypertrophic chondrocyte was defined as cells in the last lacuna that were not invaded by metaphyseal blood vessels,[5] and the number of terminal hypertrophic chondrocytes per 100 μm was defined as the column number per 100 μm width.

**Assessment of bone growth rate by alizarin injection.** To assess the rate of physical bone growth, we injected alizarin subcutaneously (30 μg/g body weight; A3883, Sigma-Aldrich) in mice at the indicated time points. Mice were euthanized 48 h after injection, and undecalcified sagittal knee sections (5 μm in thickness) were prepared. Longitudinal bone growth was evaluated as the distance between the edge of the red fluorescence-labeled metaphysis and the chondro-osseous junction.[5] The distances at four equally spaced points in a center area of the proximal tibial growth plate with a width of 600 μm were measured per section.

**Immunohistochemistry.** For p-Akt and p-H3 stainings, decalcified sagittal knee sections (5 μm in thickness) were prepared and post-fixed in 4% PFA for 5 min at room temperature. Antigen retrieval was performed by incubating the sections with 0.1% trypsin for 10 min at room temperature. After blocking with 5% goat serum (G9023, Sigma-Aldrich) in PBS for 30 min, sections were incubated with rabbit anti-p-Akt antibody (1:50; 9271S, Cell Signaling Technology) or rabbit anti-p-H3 antibody (1:100; 04-817, Sigma-Aldrich) for 60 min at room temperature, washed with PBS, and subsequently incubated with Alexa Fluor 568-conjugated goat anti-rabbit IgG (1:250; A11011, Thermo Fisher) for 60 min at room temperature. The sections were examined under 40x magnification for p-H3 and 60x magnification for p-Akt. A central area of the proximal tibial growth plate with a width of 600 μm was selected, and the ratio of p-H3$^+$ cells or p-Akt$^+$ cells among DAPI$^+$ or ZsGreen$^+$ cells was analyzed using Image J. For Igf-1 immunostaining, decalcified sagittal knee paraffin sections (6 μm in thickness) were prepared. Antigen retrieval was performed by incubating the sections with 10 mM sodium citrate (pH 6.0) for 10 min at 80C°. After blocking with 5% goat serum (G9023, Sigma-Aldrich) in PBS for 30 min, sections were incubated with rabbit anti-IGF1 antibody (1:100; ab9572, Abcam) for overnight at 4C°, washed with PBS, and subsequently incubated with Alexa Fluor 649-conjugated goat anti-rabbit IgG (1:200; A11011, Thermo Fisher) for 60 min at room temperature.

***In situ* hybridization.** RNA *in situ* hybridization was performed on proximal tibial growth plates as described previously.[6] Briefly, undecalcified sagittal knee sections (5 μm in thickness) were post-fixed in freshly prepared 2% formaldehyde for 30 min at room temperature, dehydrated with 70% ethanol for at least one day at 4 °C. The sections on the film were then placed on glass slides. Once the ethanol volatilized, the four sides of the film were surrounded with vacuum grease dissolved in spectroscopic-grade chloroform (C298-500, Sigma-Aldrich) to fix the sections onto glass slides. After the grease was dried, RNA *in situ* hybridization was performed using RNAscope 2.5HD detection reagent kit-RED (Advanced Cell Diagnostics Inc., Newark, CA, USA) according to manufacturer instructions. Sequences of the probes used are as follows (NCBI accession numbers): *Ki67* (NM_001081117.2, 1472–2523), *Cd73* (NM_011851.4,1365–2786), *Pthrp* (NM_008970.4, 173–1231), *Igf-1* (NM_010512.4, 359–1354), *Clu* (NM_013492.2, 575–1794), *Igf1r* (Custom probe), *Matn1* (NM_010769.2, 857-1795), *Col2a1* (NM_001113515.2, 711-2052) and *Slc2a1* (NM_011400.3, 325-1331). A central area of the proximal tibial growth plate with a width of 600 μm was used for analysis. We performed quantification according to the semi-quantitative scoring guideline from the manufacturer: Score 0, no staining or <1 dot/10 cells; Score 1, 1–3 dots/cell; Score 2, 4–9 dots/cell, no or very few dot clusters; Score 3, 10–15 dots/cell and/or <10% of dots in clusters; Score 4, > 15 dots/cell and/or >10% of dots in clusters (https://acdbio.com/dataanalysisguide). For genes with low expression levels, such as *Ki67*, *Cd73*, and *Pthrp*, the sections were examined under 40x magnification and the cells with at least one dot (Score 1–4) were counted as positive. For genes with relatively high expression levels, such as *Igf-1* and *Clu*, the sections were examined under a 20x magnification and the cells with more than 10 dots or with clustering dots (Score 3, 4) were counted as positive.

**EdU labelling and detection.** To evaluate cell proliferation, EdU (50 μg/g body weight, 900584, Sigma-Aldrich) dissolved in saline was subcutaneously administered to the mice at the indicated postnatal days. After incubating samples with 0.5% Triton X-100 (9002-93-1, Sigma-Aldrich) in PBS for 10 min at room temperature, the Click-iT EdU Alexa Fluor 647 Imaging Kit (C10340, Thermo Fisher) was used to detect

EdU. A central area of the proximal tibial growth plate with a 600 μm width was used for analysis. The ratio of the number of $EdU^+$ cells among $DAPI^+$ cells was measured by BZ-X Analyzer.

**Measurement of serum IGF-1 levels.** Serum IGF-1 levels were determined using a mouse-specific Quantikine ELISA kit (R&D Systems, MG100) according to manufacturer protocol.

**LMD and RNA-Seq analysis.** We performed LMD using snap frozen sections of distal femur and proximal tibial growth plates as previously described.[7] Briefly, an adhesive film was attached to the specimen and 8 μm sections were cut. Frozen specimens were immediately soaked in 80% ethanol for 30 s, then in 100% ethanol for 1 min (twice), then in xylene for 5 min at room temperature, and subsequently dried for 5 min. Because most *Axin2-CreER*$^+$ cells remained morphologically in the resting zone after seven days of chase (Fig. 2*B*), we utilized *Axin2Cre*$^{ERT2}$*;R26R* reporter system as a tool to label the resting chondrocytes. We used *Axin2Cre*$^{ERT2}$*;R26R*$^{TdTomato}$ mice instead of *Axin2Cre*$^{ERT2}$*;R26R*$^{ZsGreen}$ mice because ZsGreen signal was diminished during ethanol fixation.[8] Knee samples were prepared from *Axin2Cre*$^{ERT2}$*;R26R*$^{TdTomato}$ mice on P30 which received tamoxifen injections for three consecutive days from P23 (Fig. S5a). Microdissections of TdTomato-positive resting chondrocytes and unlabeled proliferative chondrocytes were performed using a Leica LMD 7000 laser microdissection system (Leica Microsystems). We defined proliferative chondrocytes by flattening pattern and location at least 50 μm from the SOC. RNA extraction was performed using the Picopure RNA Isolation Kit (KIT0204, Thermo Fisher) and was amplified by two rounds of *in vitro* transcription using the Arcturus RiboAmp HS PLUS Kit (KIT0521, Thermo Fisher), according to manufacturer protocol. Complimentary DNA libraries were prepared using the KAPA RNA HyperPrep Kit (Roche, Basel, Switzerland) according to manufacturer instructions. The unique dual barcode sequences were incorporated into adaptors for multiplexed high-throughput sequencing (NEXTFLEX® Unique Dual Index Barcodes; BioO Scientific, Austin, TX, USA). The libraries were pooled and loaded onto an S1 flow cell on an Illumina NovaSeq 6000 (Illumina, San Diego, CA, USA) and run for 57-65 cycles. De-multiplexed and adapter-trimmed sequencing reads were generated using Dragen

Host Software Version 01.011.588.3.7.5 (Illumina), allowing no mismatches in the index read. BBDuk (version 36_49) was used to trim/filter low quality sequences using the "qtrim=lr trimq=10 maq=10" option. Next, the filtered were aligned to the mouse reference genome (GRCm38) using HISAT2 (version 2.1.0) by applying the -no-mixed and -no-discordant options. The reads were summarized for each gene using HTseq-count (version 0.11.2) supplemented with Ensembl gene annotation (GRCm38.78). The edgeR R package was used to fit the read counts to a negative binomial model along with the generalized linear model, and differentially expressed genes were determined by the likelihood ratio test method in edgeR. Significance was defined to be those with q-value < 0.01 calculated by the Benjamini-Hochberg method to control the false discovery rate (FDR) and log2 fold change is greater than 1 or smaller than -1 in resting chondrocytes compared to proliferative chondrocytes. The ggpubr R package was used to generate a MA plot. The top 500 DEGs were selected, and the ComplexHeatmap R package was used to generate a heatmap. The list of DEGs was subjected to KEGG pathway analysis using the Database for Annotation, Visualization, and Integrated Discovery (DAVID) bioinformatics resource (david.ncifcrf.gov). We also used the Upstream Regulator Analysis tool in Ingenuity Pathways Analysis Software (version 01-20-04) (Ingenuity Systems Inc., Redwood, CA, USA) to predict the potential cause of the changes in gene expression between resting and proliferative chondrocytes.

**Chondrocyte isolation and culture.** Chondrocytes were isolated from the growth plate following the protocol previously described with some modifications.[9] Briefly, the femur and tibia bones were incubated in calcium- and magnesium-free Hank's Balanced Salt Solution (HBSS; H6648, Sigma) containing 2 units of liberase TM (Roche) at 37°C for 30 minutes. After removing soft tissues including perichondrium, distal femur, and proximal tibia bones were dislodged from the the diaphyseal end at the boundary between the primary spongiosa and the growth plate. These epiphyseal bones containing the growth plate were then digested in HBSS with 2 units of liberase TM at 37°C for 30 minutes on a rotor. The first digestion solution was collected and centrifuged at 500 g at 4°C for 4 minutes to collect cells followed by additional four sequential digestion steps at 37°C for 30 minutes on a rotor. The harvested cells were pooled for cell culture and flow cytometry analysis.

The isolated chondrocytes were seeded at the density of 3.4 x $10^4/cm^2$ and passaged at the split ratio of 1:4 twice. At Passage 3, some cells were seeded at the density of 4.2 x $10^3/cm^2$ and serially passage until they stopped growing. The number of cells were counted at each passage (n=3). At Passage 3, the rest cells were sorted by the flow cytometry. The ZsGreen positive cells were collected and cultured under chondrogenic, osteogenic and adipogenic differentiation conditions: In chondrogenic culture, the cells were seeded at the density of 12.5 x$10^4/cm^2$ in 96-well plate, cultured in high-glucose DMEM containing 10% FBS and 50 μg/ml ascorbic acid for 7 days; In osteogenic differentiation culture, the cells were seeded at 4.2x$10^3/cm^2$ and cultured in StemXVivo Osteogenic/Adipogenic BaseMedia (R&D systems, Inc., Minneapolis, MN) with Mouse/Rat Ostegenic Supplement (R&D systems, Inc.) for 7 days. In adipogenic differentiation culture, the cells were seeded at 2.1 x $10^4/cm^2$ and cultured in was in StemXVivo Osteogenic/Adipogenic BaseMedia with Human/Mouse/Rat Adipogenic Supplement (R&D systems, Inc.) for 14 days. At the end of culture, the cultures were stained with alcin blue (pH 1.0), alizarine red S (pH 4.1) or Oil red O.

**Flow cytometry analysis.** The chondrocytes were subjected to flow cytometry (Cytek® Aurora, 4 Laser 16UV-16V-14B-8R) and the obtained data was analyzed by FlowJo 10.8.1 (BD Biosciences). The following antibodies were used: anti-CD45 (30-F11, APC-conjugated, 1:200, BioLegend), anti-Ter119 (TER-119, APC-conjugated, 1:200, BioLegend), anti-Tie2 (TEK4, APC-conjugated, 1:200, BioLegend), anti-CD51 (RMV-7, biotinylated, 1:100, BioLegend), anti-Thy1.1 (OX-7, BV421-conjugated, 1:100, BioLegend), anti-Thy1.2 (30-H12, BV421-conjugated, 1:100, BioLegend), anti-6C3 (BP-1, BV711-conjugated, 1:100, BD Biosciences), anti-CD105 (MJ7/18, PE-Cy7-conjugated, 1:100, BioLegend), anti-CD200 (OX-90, APC-R700-conjugated, 1:100, BD Biosciences), APC/Cy7 Streptavidin (1:200, BioLegend). DAPI was used to exclude dead cells.

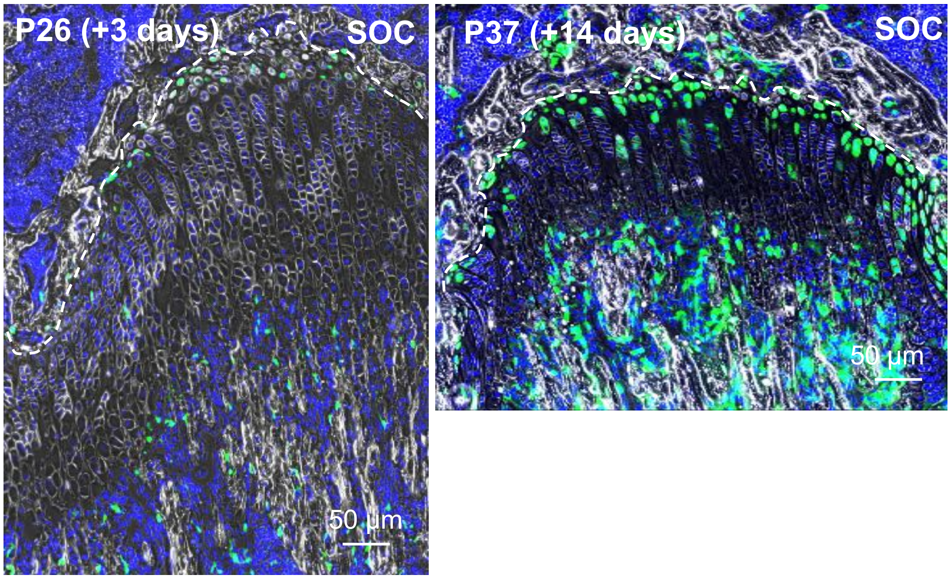


**Supplementary Figure S1.** Fate-mapping analysis of Axin2$^+$ cells in the distal femoral growth plate in *Axin2Cre$^{ERT2}$;R26R$^{ZsGreen}$* mice (pulsed on P23–25, and traced for 3 and 14 days). Representative images showing descendants of initially labeled Axin2$^+$ cells at 3 (n=3) (P26, + 3 days) and 14 (n=3) (P37, +14 days) days after tamoxifen injection. Axin2+ cells were detected at the border of secondary ossification center (SOC) in the resting zone of the distal femoral growth plate 3 days after tamoxifen injection. The Axin2+ cells increased in the resting zone and made columns in the growth plate 14 days after tamoxifen injection. The white dashed lines demarcate the growth plate from the SOC. Scale bar:50 µm.

**a** Fractionation by SSC markers

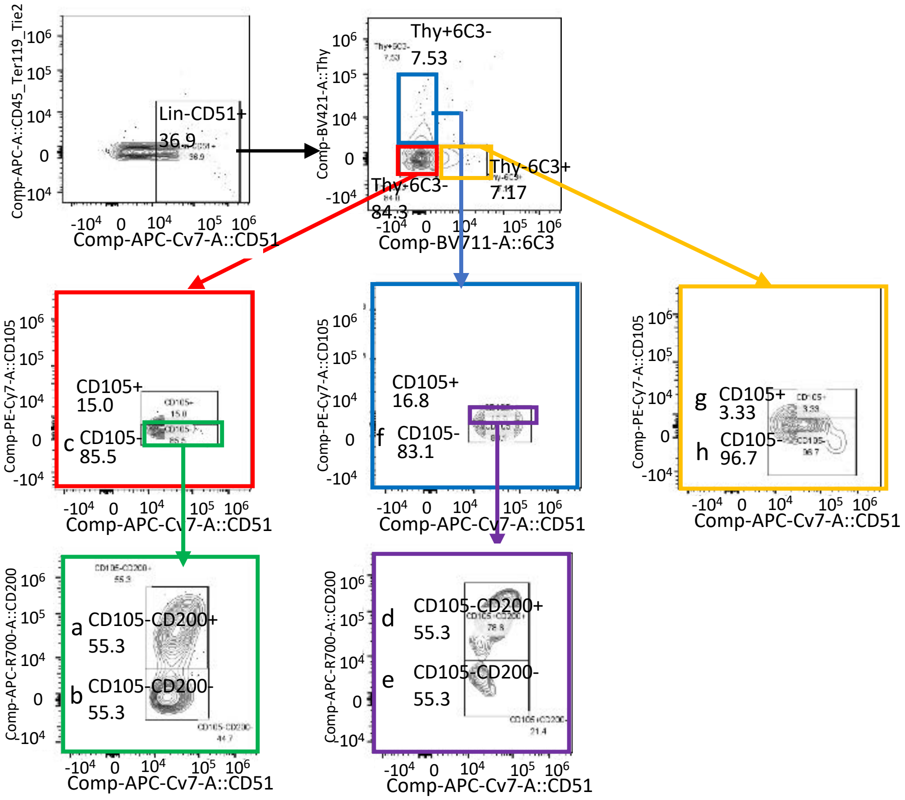

**b** The ratio of Axin2+ cells

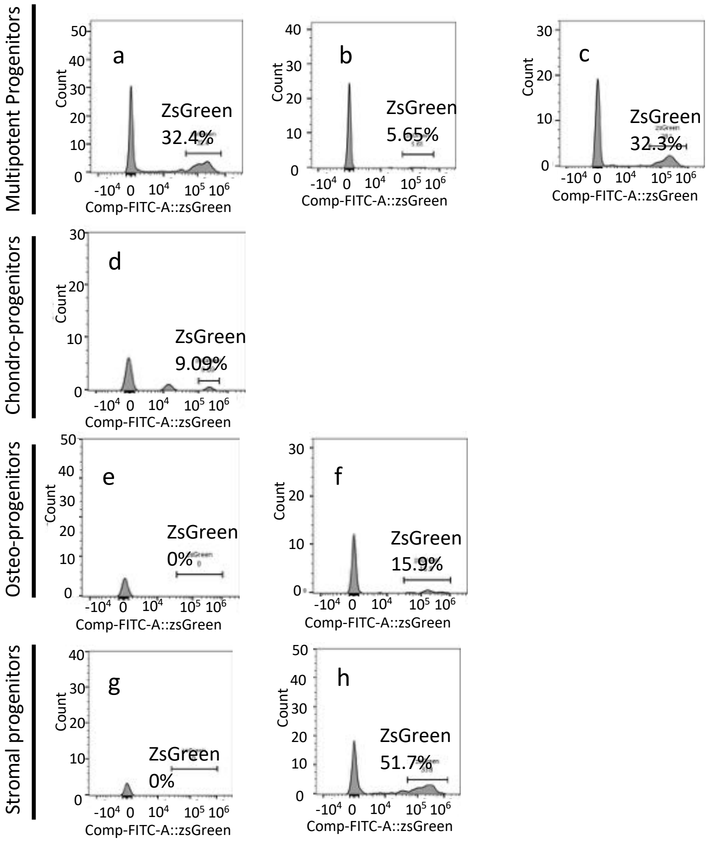


**c** Expression of SSC markers in Axin2+ cells

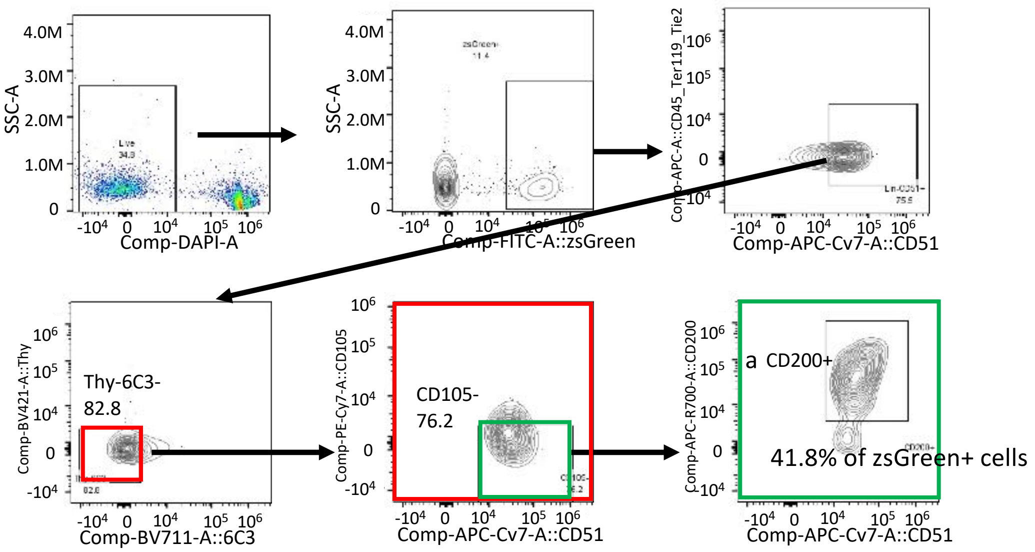

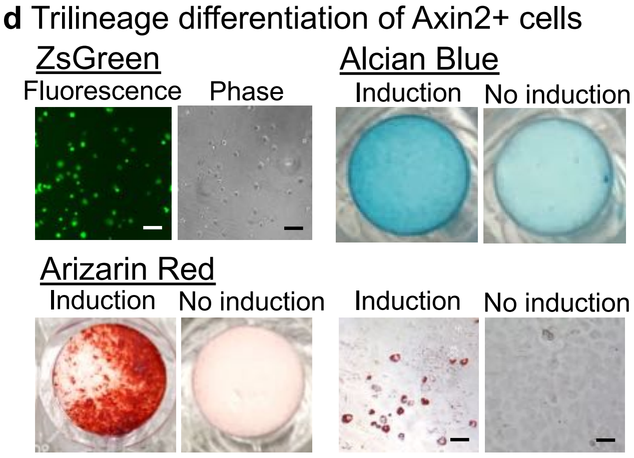


**Supplementary Figure S2.** Characterization of Axin2+ cells and their descendants. The growth plate chondrocytes were isolated from proximal tibia and distal femur in the $Axin2Cre^{ERT2};R26R^{ZsGreen}$ mice (pulsed on P23–25, and harvested on P26 (a-c) or P37 (d)). **a** Fractionation of Axin2+ cells by skeletal stem cell markers. The growth plate cells were subjected to flow cytometry to analyze expression of skeletal stem cells (SSC) markers following the procedure described previously.[10] **b** FACS plot for fractions a-h shown in (a) by ZsGreen signal. The values show the percentage of ZsGreen-positive Axin2+ cells. **c** FACS plot of the ZsGreen+ cells. 41.8% of ZsGreen-positive Axin2+ cells were fractioned in the multipotent progenitor population (a, CD45-TER119-Tie2-AlphaV+Thy-6C3-CD105-CD200+). **d** Multipotency of Axin2+ cells. The growth plate cells were passaged twice and subjected to flow cytometry. The ZsGreen+ cells were sorted and seeded at the density of 12.5 $x10^4/cm^2$ in 96-well plate (chondrogenic differentiation), $4.2x10^3/cm^2$ in 48-well plate (osteogenic differentiation) and 2.1 x $10^4/cm^2$ in 48-well plate (adipogenic differentiation). On day 1, the fluorescence and phase contrast images were taken (ZsGreen). The cells cultured in the chondrogenic differentiation or undifferentiated medium were stained with alcian blue on day 7 (Alcian Blue). The cells cultured in the osteogenic differentiation or undifferentiated medium were stained with alcian blue on Day 7 (Alizarin Red). The cells cultured in the adipogenic differentiation or undifferentiated medium were stained with oil red O on Day 14 (Oil Red O). Scale bars: 50 μm.

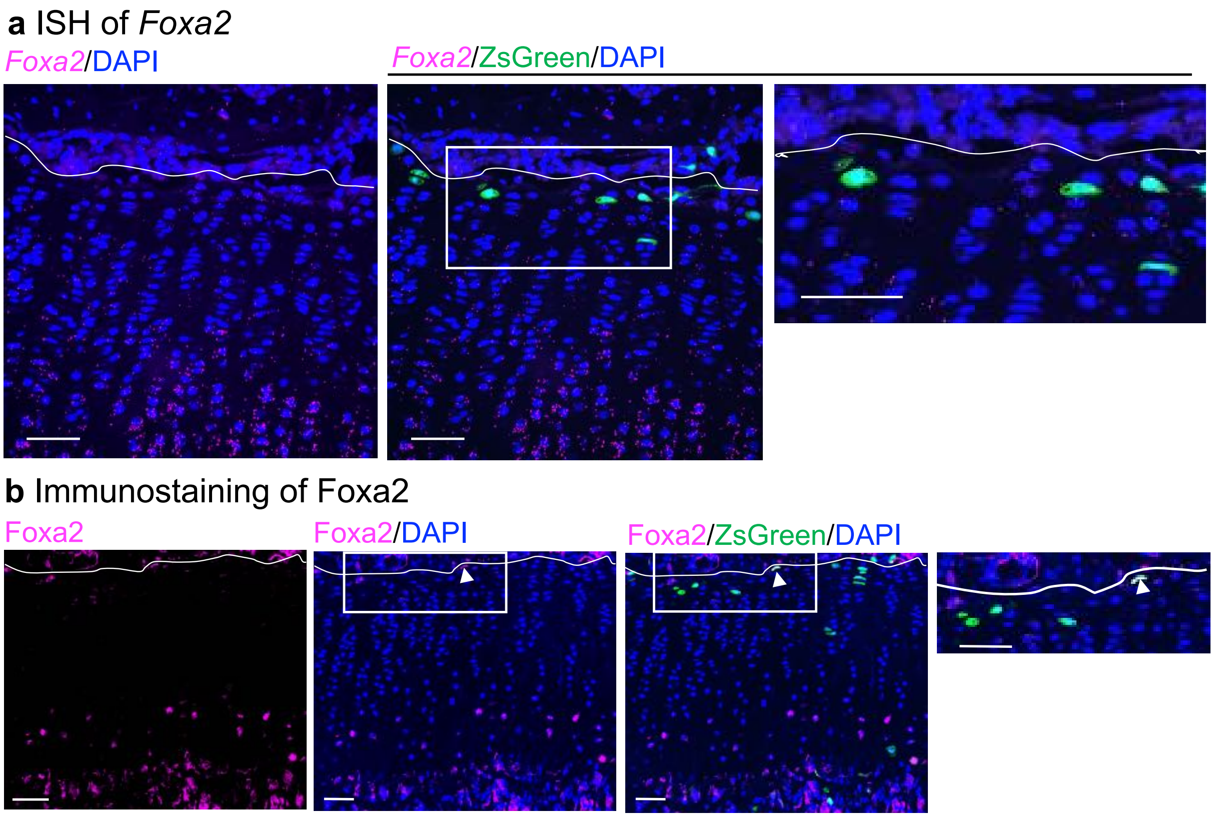


**Supplementary Figure S3. *In situ* hybridization and immunostaining of Foxa2.** The proximal tibial growth plate was collected from the *Axin2Cre$^{ERT2}$;R26R$^{ZsGreen}$* mice (pulsed on P23–25, and harvested on P26) and subjected to histological analysis. **a** Representative images of *in situ* hybridization for *Foxa2* and Axin2+ cells (n=3). The *Foxa2* transcripts were sparsely detected in the resting and proliferative zones and abundantly in the hypertrophic zone. **b** Representative images of immunostaining for Foxa2 and Axin2+ cells (n=3). Immunoreactivity for Foxa2 were detected in some resting chondrocytes and hypertrophic chondrocytes. Some Axin2+ cells were positive for Foxa2 immunostaining. The white dashed lines demarcate the growth plate from the secondary ossification center. Scale bars: 50 µm.

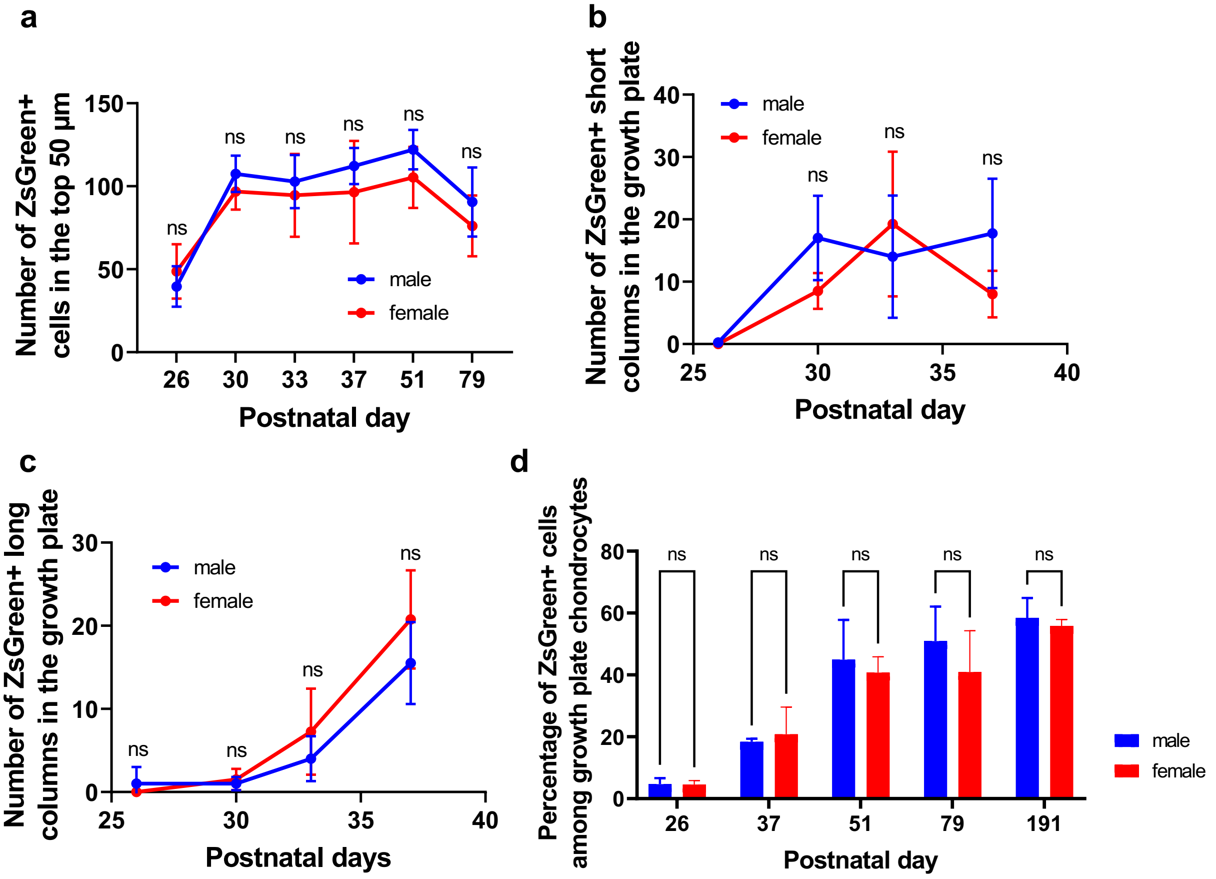


**Supplementary Figure S4. No significant difference between sex in the lineage tracing results in *Axin2Cre$^{ERT2}$;R26R$^{ZsGreen}$* mice. a–d** Fate-mapping analysis of Axin2$^{+}$ cells in the proximal tibial growth plate in *Axin2Cre$^{ERT2}$;R26R$^{ZsGreen}$* mice (pulsed on P23–25, and traced for several periods). Quantification of descendants of initially labeled Axin2$^{+}$ cells (ZsGreen$^{+}$ cells) in the top 50 μm (**a**), ZsGreen$^{+}$ columns in the growth plate, short columns (≤ 9 cells, red line) (**b**) and long columns (≥ 10 cells, blue line) (**c**), and percentage of ZsGreen$^{+}$ cells among growth plate chondrocytes (**d**). P26–51 (n = 4 male, n = 4 female), P79 (n = 4 male, n = 3 female), P191 (n = 3 male, n = 3 female). Adjusted $p > 0.9999$, (ns) (P26–79) (**a**). Adjusted $p > 0.9999$, (ns) (P26 and P33), adjusted $p = 0.3261$, (ns) (P30), adjusted $p = 0.4373$, (ns) (P37) (**b**). Adjusted $p > 0.9999$, (ns) (P26–P33), adjusted $p = 0.8919$ (ns) (P37) (**c**). Adjusted $p > 0.9999$, (ns) (P26–P191) (**d**). ns, not significant. All data are presented as the mean ± SD. Statistical significance between sex was determined by two-way analysis of variance and Bonferroni's multiple comparison test.

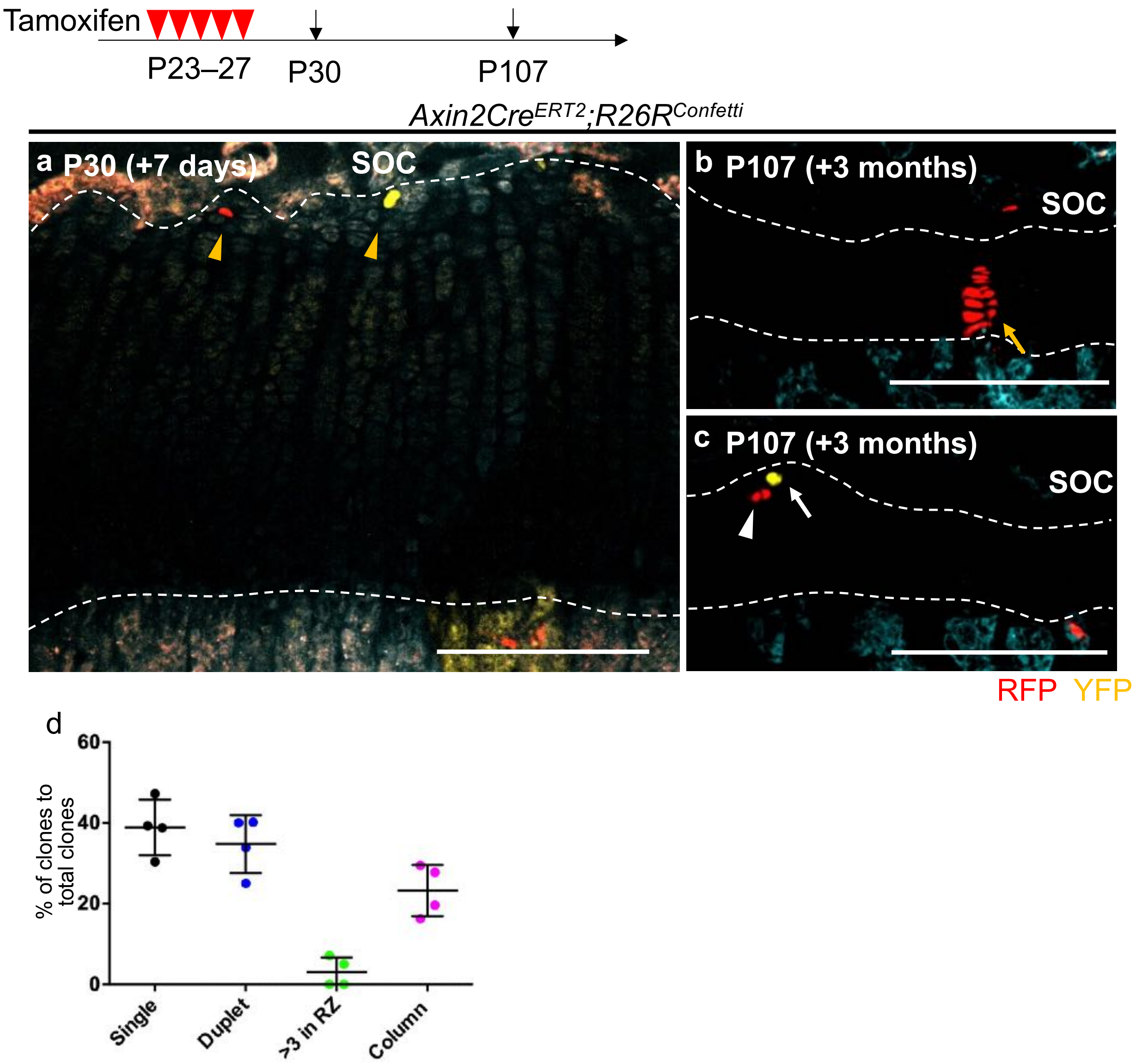


**Supplementary Figure S5. *In vivo* clonal analysis of Axin2$^{+}$ cells. a–c** Clonal fate-mapping analysis of Axin2$^{+}$ cells in the proximal tibial growth plate in *Axin2Cre$^{ERT2}$;R26R$^{Confetti}$* mice (pulsed on P23–27, and traced for 7 days or 3 months). Orange arrowheads, Axin2$^{+}$ cells (**a**), orange arrow, chondrocyte column in the same color (**b**), white arrow, singlet progeny, white arrowhead, doublet progeny (**c**). **d** Quantification of the percentage of single-cell clones (black), duplet clones (blue), the clones of more than 3 cells (green) in the resting zone and the clones making columns in the growth plate (magenta) to the total number of clones. The data are presented as the mean ± SD. SOC, secondary ossification center. The white dashed lines demarcate the growth plate from the surrounding tissues. Scale bars: 200 μm (**a**–**c**).

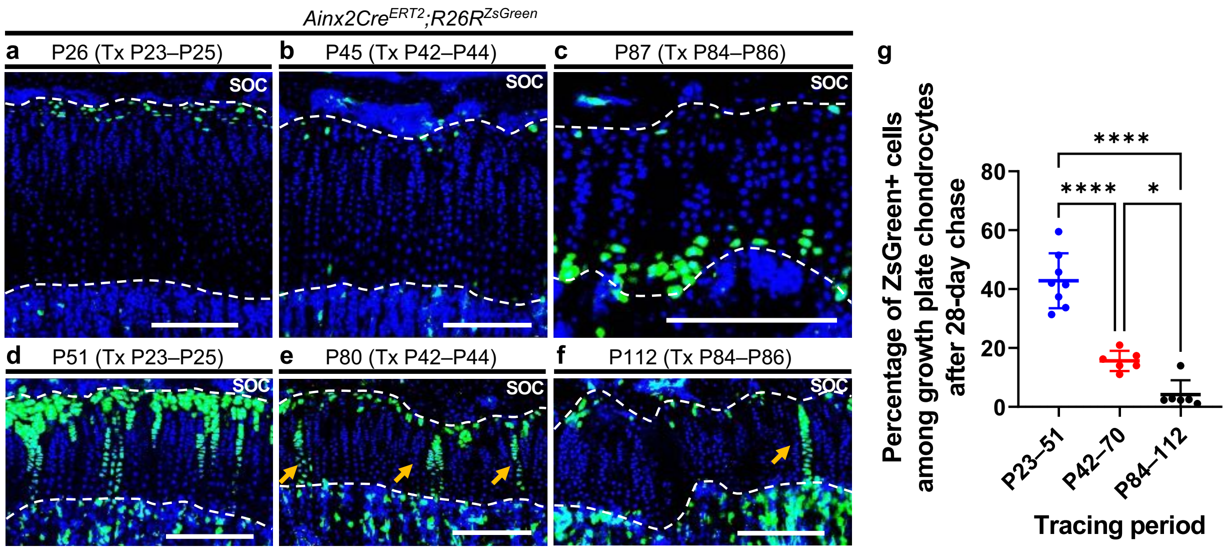


**Supplementary Figure S6. Chondrocyte column formation of Axin2+ cells at different developmental stages. a**–**f** Fate-mapping analysis of Axin2$^{+}$ cells in the proximal tibial growth plate in *Axin2Cre$^{ERT2}$;R26R$^{ZsGreen}$* mice (pulsed for three consecutive days from various time points and chased for 3 or 28 days). Axin2$^{+}$ cells were labelled with tamoxifen injection from P23 (**a**), P42 (**b**), and P84 (**c**). Representative images of the descendants of initially labeled Axin2$^{+}$ cells (ZsGreen$^{+}$ cells) chased for 28 days from P23 (**d**), P42 (**e**), and P84 (**f**). Arrows, ZsGreen$^{+}$ chondrocyte columns (**e**, **f**). **g** Percentage of ZsGreen$^{+}$ cells among growth plate chondrocytes after a 28-day chase at various time points. P23–51 (n = 8), P42–70 (n = 6), P84–112 (n = 6). Adjusted ****p= < 0.0001 (P23–51 vs P42–70, P23–51 vs P84–112), adjusted *p = 0.0245 (P42–70 vs P84–112). Tx, tamoxifen, SOC, secondary ossification center. The white dashed lines demarcate the growth plate from the surrounding tissues. Scale bars: 200 µm (**a**–**f**). All data were presented as the mean ± SD. Statistical significance was determined by one-way analysis of variance and Tukey's multiple comparison test.

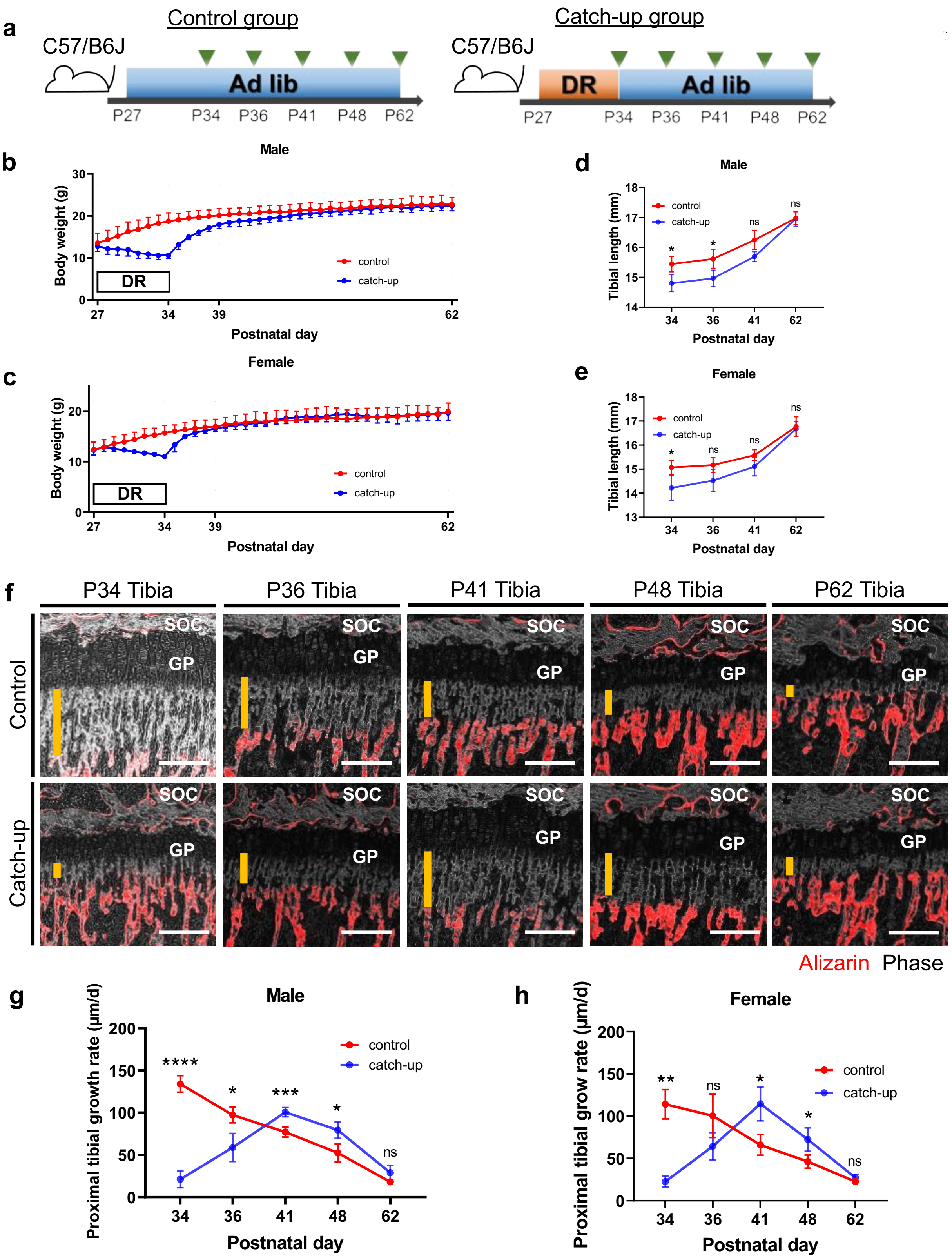
a
Control group
C57/B6J
Ad lib
P27
P34
P36
P41
P48
P62
Catch-up group
C57/B6J
DR
Ad lib
P27
P34
P36
P41
P48
P62
b
Male
Body weight (g)
DR
control
catch-up
Postnatal day
c
Female
Body weight (g)
DR
control
catch-up
Postnatal day
d
Male
Tibial length (mm)
control
catch-up
Postnatal day
e
Female
Tibial length (mm)
control
catch-up
Postnatal day
f
P34 Tibia
P36 Tibia
P41 Tibia
P48 Tibia
P62 Tibia
Control
Catch-up
SOC
GP
Alizarin Phase
g
Male
Proximal tibial growth rate (μm/d)
control
catch-up
Postnatal day
h
Female
Proximal tibial grow rate (μm/d)
control
catch-up
Postnatal day

**h** Growth plate analysis in Control and DR

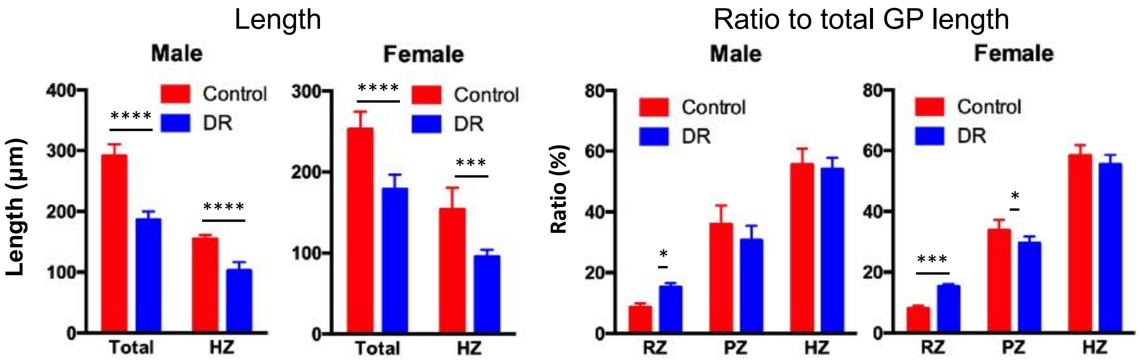


**i** Growth plate analysis in Control and Catch-up

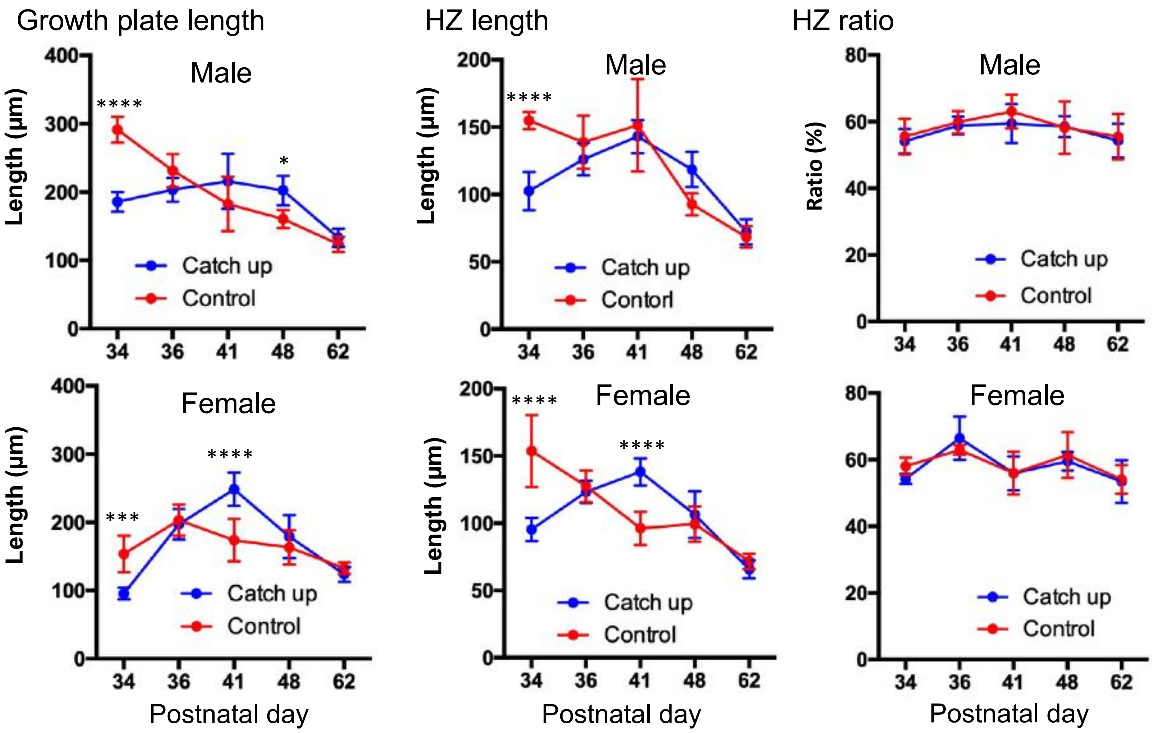


**j** Proliferation

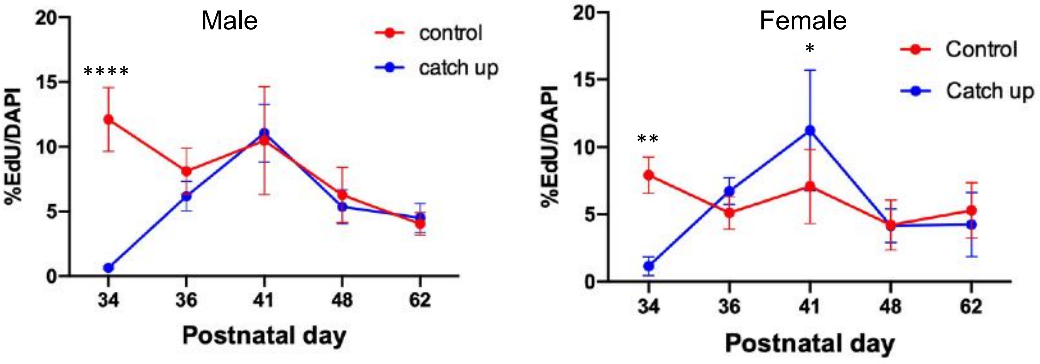

**Supplementary Figure S7. Catch-up growth during ad libitum feeding after a seven-day dietary restriction in C57/B6j mice. a** Schematic diagrams of the mouse catch-up growth model examining the proximal tibial growth plate of C57/B6J mice. The mice were fed ad libitum from P27 (control group) or after seven-day DR from P27 (catch-up group). **b**, **c** Weight changes during and after DR in male (**b**) and female mice (**c**). n = 5 (each time point) for male and female mice. **d**, **e** Tibial length changes after stopping DR in male (**d**) and female mice (**e**). P34 (n = 6 control, n = 5 catch-up), P36–62 (n = 5 control, n = 5 catch-up) in male mice (**d**), P34 (n = 7 control, n = 7 catch-up), P36, (n = 7 control, n = 6 catch-up), P41, P62 (n =5 control, n = 5 catch-up) in female mice (**e**). Adjusted *p = 0.0186 (P34), adjusted *p = 0.00331 (P36), adjusted p = 0.0562 (ns) (P41), adjusted p > 0.9999 (ns) (P62) (**d**). Adjusted *p = 0.0163 (P34), adjusted p = 0.0714 (ns) (P36), p = 0.2246 (ns) (P41), p > 0.9999 (ns) (P62) (**e**). **f** Representative images for alizarin labelling in the proximal tibial growth plate after DR. Mice were subcutaneously injected with alizarin (30 µg/g body weight) and euthanized after 48 h. Orange bars indicate bone growth after 48 h. **g** Proximal tibial growth rate after stopping DR in male and female mice. **h** Lengths of total growth plate and hypertrophic zone (HZ), and ratios of resting zone (RZ), proliferative zone (PZ) and hypertrophic zone (HZ) lengths to total growth plate length in control and DR groups at P34. **i** Lengths of total growth plate and HZ, and ratios of HZ length to total growth plate length in control and catch up growth groups at P34-P62 after stopping DR. **j** Percentage of EdU+ cells to DAPI + cells in the growth plate in control and catch-up growth groups on P34-P62 after stopping DR. P34 (n = 6 control, n = 5 DR), P36–P62 (n = 5 control, n = 5 catch-up) in male mice (**g-i**), P34 (n = 5 control, n = 5 catch-up), P36 (n = 5 control, n = 5 catch-up), P41–P62 (n = 5 control, n = 5 control) in female mice (**g-i**). P34 (n = 3 control, n = 3 catch-up), P36–P62 (n = 5 control, n = 5 catch-up) in male and female mice (**k**). Adjusted ****p < 0.0001 (P34), adjusted *p = 0.0180 (P36), adjusted ***p = 0.0010 (P41), adjusted *p = 0.0164 (P48), adjusted p = 0.2096 (ns) (P62) in male (**g**). Adjusted **p = 0.0039 (P34), adjusted p = 0.1939 (ns) (P36), adjusted *p = 0.0138 (P41), adjusted *p = 0.0483 (P48), adjusted p = 0.3939 (ns) (P62) in female (**g**). Adjusted ****p<0.0001 (length), adjusted***p=0.0001 (length), adjusted*p=0.0481 (ratio, male), adjusted***p=0.0001 (ratio, female), adjusted*p=0.0291 (ratio, female) (**h**). Adjusted****p<0.0001, adjusted*p=0.0361 (growth plate) in male mice, Adjusted****p<0.0001m adjusted***p=0.0003 in female mice (**i**). Adjusted ****p<0.0001, adjusted**p=0.0040, *p=0.0305 (**k**). Ad lib, ad libitum, DR, dietary restriction, ns, not significant, SOC, secondary ossification center, GP, growth plate. Scale bar; 200 µm (**f**). All data were presented as the mean ± SD. Statistical significance was determined using two-way analysis of variance and Bonferroni's multiple comparison test.

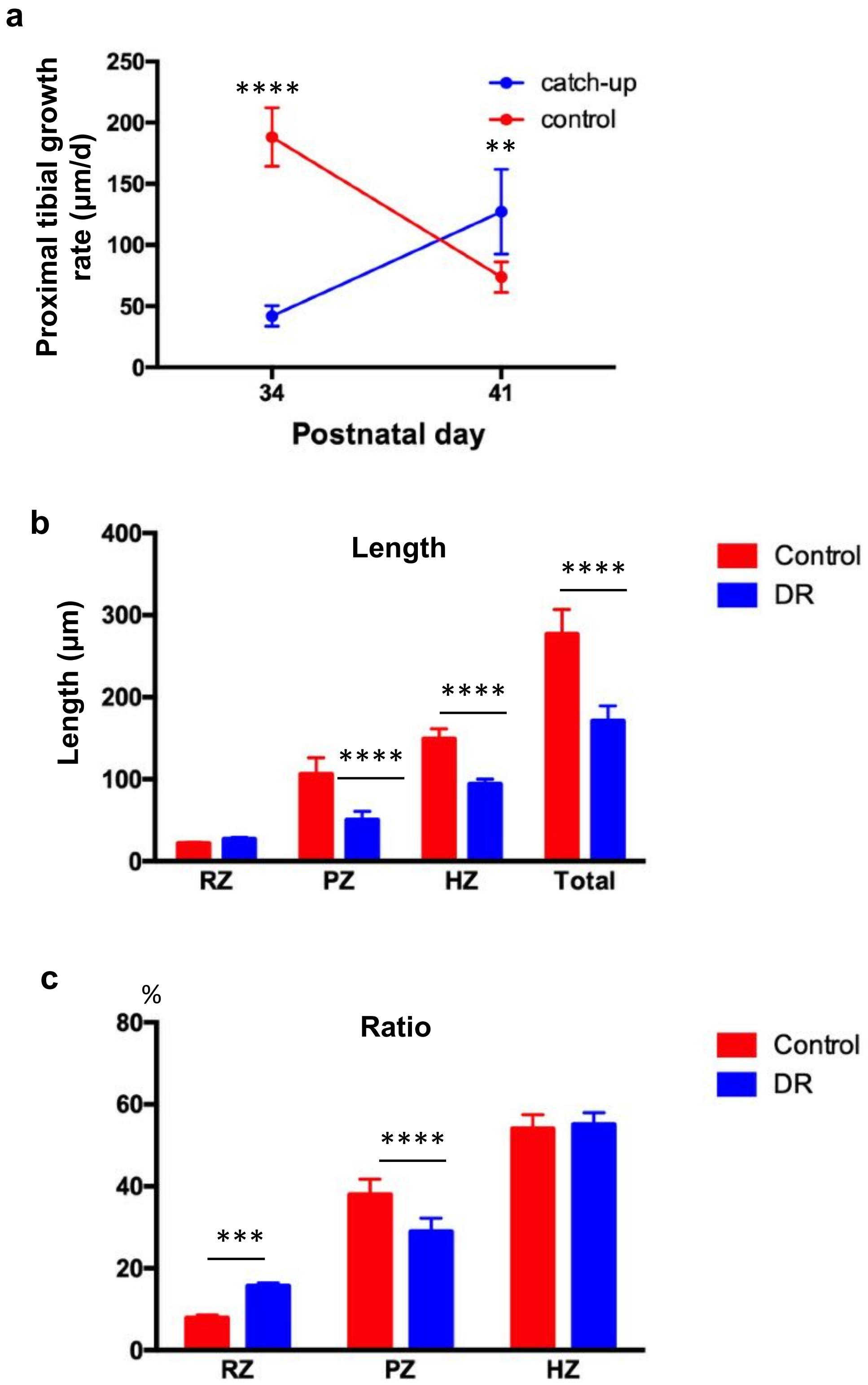
a
Proximal tibial growth rate (µm/d)
250
200
150
100
50
0
34
41
Postnatal day
catch-up
control
****
**
b
Length
Length (µm)
400
300
200
100
0
RZ
PZ
HZ
Total
Control
DR
****
****
****
c
Ratio
%
80
60
40
20
0
RZ
PZ
HZ
Control
DR
***
****

**Supplementary Figure S8. Catch-up growth during ad libitum feeding after a seven-day dietary restriction in *Axin2Cre$^{ERT2}$;R26R$^{ZsGreen}$* mice.** The *Axin2Cre$^{ERT2}$;R26R$^{ZsGreen}$* female mice (pulsed on P23–25) were fed ad libitum for 7 days from P27 (control group) or after seven-day DR from P27 (catch-up group)**.** Mice were subcutaneously injected with alizarin (30 μg/g body weight) 48h prior to euthanization. P34 (n=5 control, n=5 catch-up); P41 (n=5 control, n=5 catch-up). **a** Proximal tibial growth rate after stopping DR in female. Adjusted ****p <0.0001, adjusted **p=0.0023. **b, c** Lengths of total growth plate, resting zone (RZ), proliferative zone (PZ) and hypertrophic zone (HZ), and ratios of RZ, PZ and HZ lengths to total growth plate length in control and catch-up growth groups. Adjusted ****p <0.0001 (**b**). Adjusted ****p <0.0001, adjusted ***p=0.0002 (**c**). All data were presented as the mean ± SD. Statistical significance was determined using two-way analysis of variance and Bonferroni's multiple comparison test.

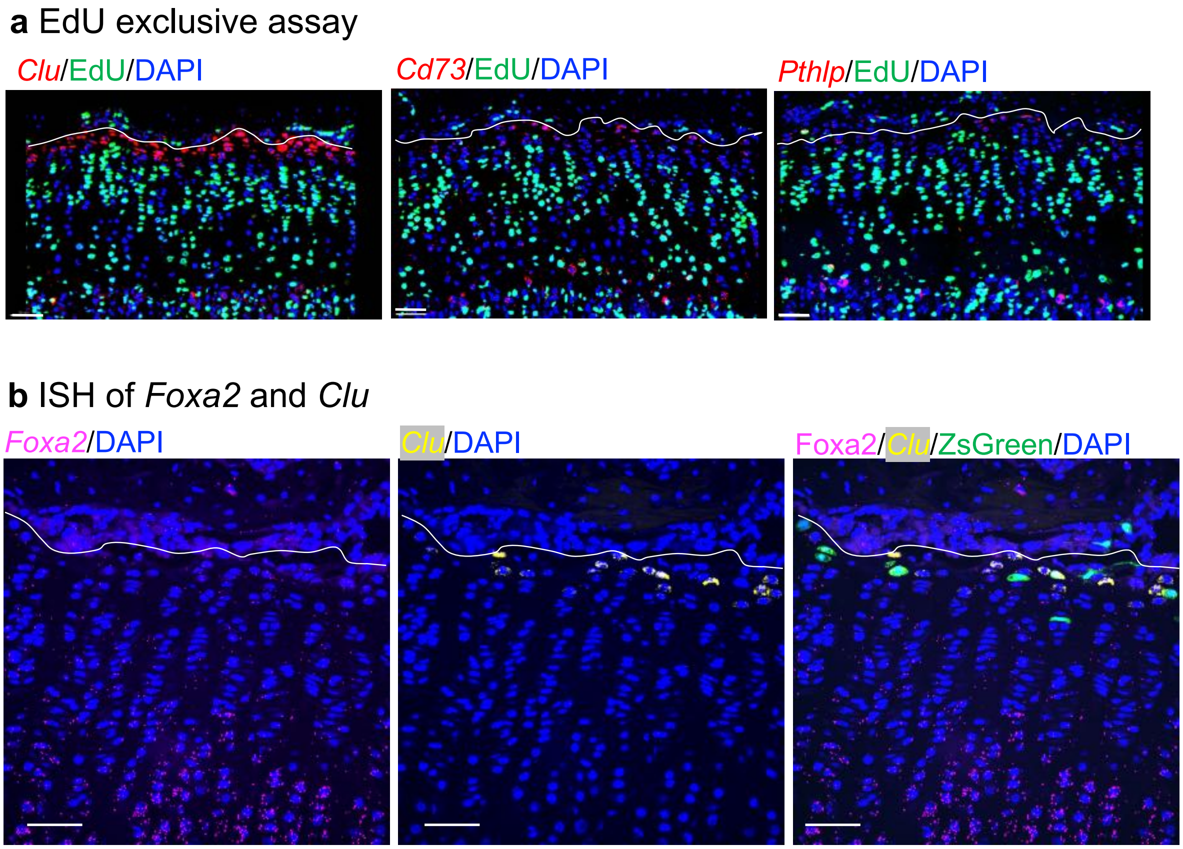


**Supplementary Figure S9**. Expression of chondroprogenitor markers. **a** Representative images for *in situ* hybridization of *Clu*, *Cd73* and *Pthlp* (red) and EdU labeling (green) in the proximal tibial growth plate (n=2). The C57/B6J mice received four consecutive daily injections of EdU (50 µg/g body weight) from P23. Two hours after the last Edu injection, the hind limbs were collected. **b** Representative images for in situ hybridization of *Foxa2* (magenta) and *Clu* (yellow), and Axin2+ cells (green) in the proximal tibial growth plate in the *Axin2Cre*$^{ERT2}$*;R26R*$^{ZsGreen}$ mice (pulsed on P23-P25 (n=3). At P26, the hind limbs were collected. The white dashed lines demarcate the growth plate from the secondary ossification center. Scale bars: 50 µm.

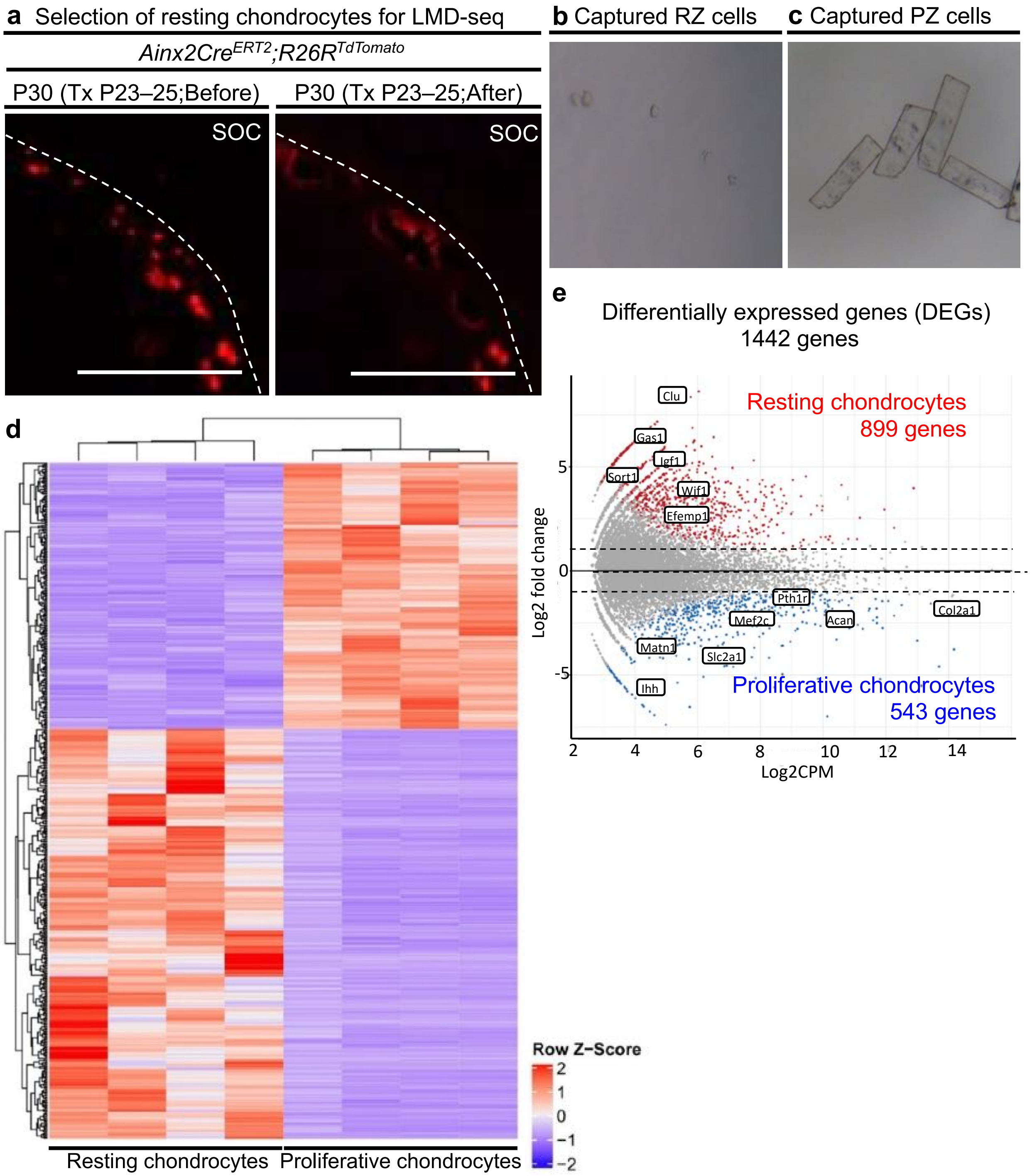
a Selection of resting chondrocytes for LMD-seq
Ainx2CreERT2;R26RTdTomato
P30 (Tx P23–25;Before)
P30 (Tx P23–25;After)
SOC
SOC
b Captured RZ cells
c Captured PZ cells
d
Resting chondrocytes
Proliferative chondrocytes
Row Z-Score
2
1
0
−1
−2
e
Differentially expressed genes (DEGs)
1442 genes
Resting chondrocytes
899 genes
Proliferative chondrocytes
543 genes
Clu
Gas1
Igf1
Sort1
Wif1
Efemp1
Pth1r
Col2a1
Mef2c
Acan
Matn1
Slc2a1
Ihh
Log2 fold change
Log2CPM
5
0
-5
2
4
6
8
10
12
14

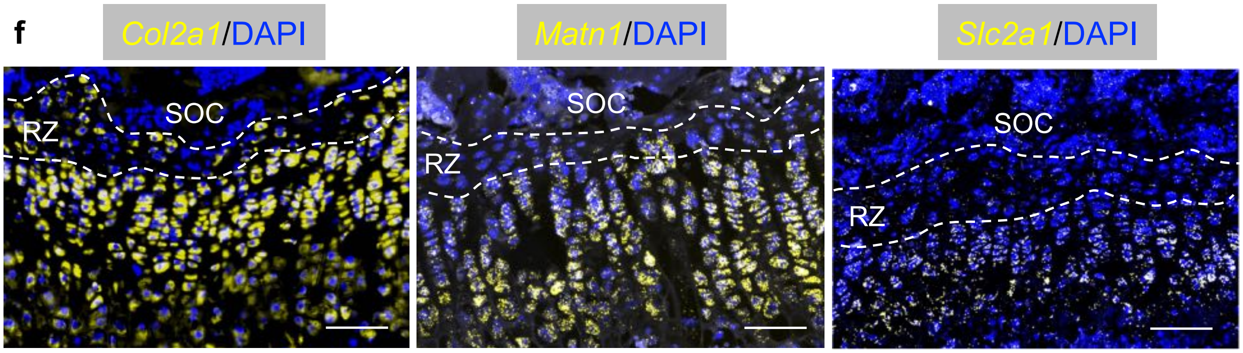


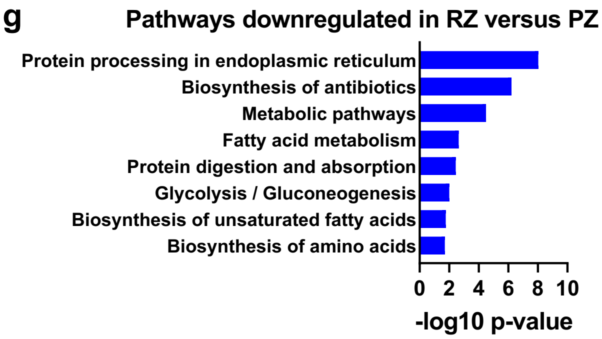


**Supplementary Figure S10. Changes in the transcriptome profiles of resting and proliferative chondrocytes analyzed by RNA-seq from laser microdissected samples. a–c** *Axin2Cre*$^{ERT2}$*;R26R*$^{TdTomato}$ mice received three consecutive daily injections of tamoxifen from P23 and samples were collected at P30 when the resting chondrocytes were labeled with red fluorochrome. (**a**) Resting chondrocytes from the P30 distal femoral growth plate were imaged before and after captured by LMD. The white dashed lines demarcate the growth plate from the surrounding tissues. Scale bars: 200 μm. **b**, **c** Captured resting chondrocytes (**b**) and proliferative chondrocytes (**c**). **d** Comparative RNA-seq analysis of resting chondrocytes and proliferative chondrocytes. Heat map of the top 500 DEGs (fold change > ± 2, FDR < 0.01) with hierarchical clustering. **e** MA plot of DEGs between resting chondrocytes (899 genes) and proliferative chondrocytes (543 genes) with representative upregulated genes in each cell population. **f** Representative images of *in situ* hybridization (yellow) of *Col2a1*, *Matn1* and *Slc2a1* (Glut1) in the proximal tibial growth plate of 4 weeks-old C57/B6J mice. **g** KEGG pathways enriched by genes significantly downregulated in the resting zone compared to the proliferative zone. Logarithmic p-values of significance are indicated on the *x*-axis. SOC, secondary ossification center. DEG, differentially expressed gene, KEGG, Kyoto Encyclopedia of Genes and Genomes, LMD, laser microdissection, RZ, resting zone, PZ, proliferative zone. The white dashed lines demarcate the resting zone (RZ) from the proliferative zone. Scale Bars: 50 µm.

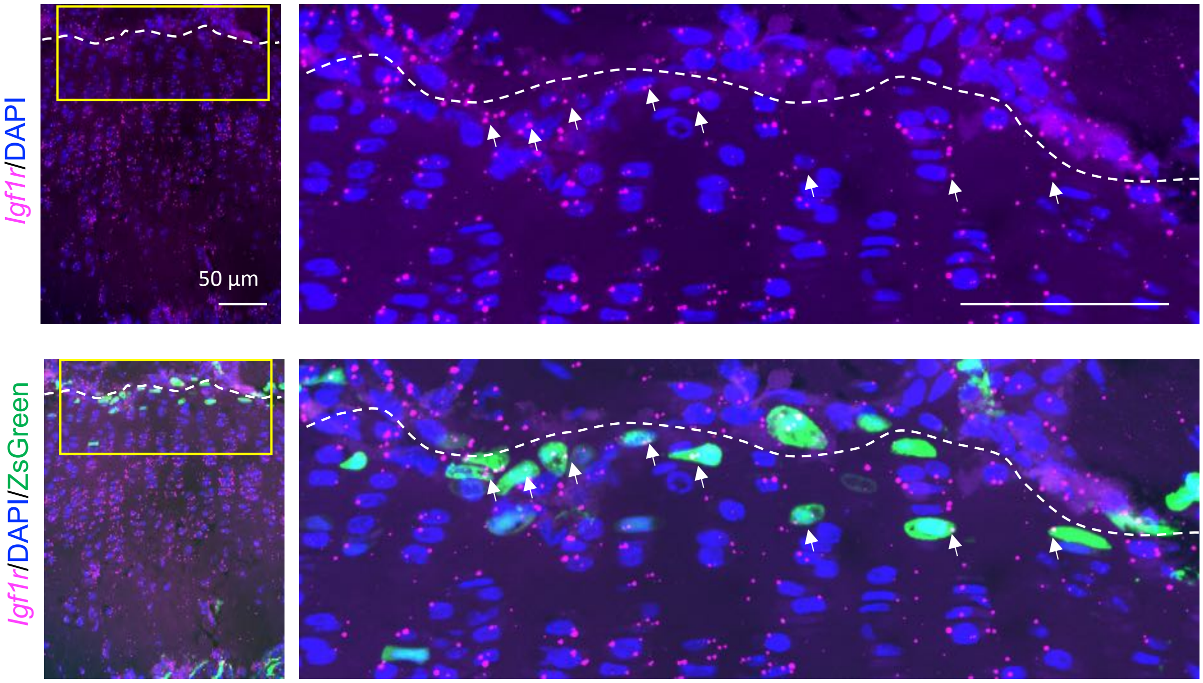


**Supplementary Figure S11. Expression of *Igf1r* in the proximal tibial growth plate.** Representative images for *in situ* hybridization of *Igf1r* (magenta) and Axin2+ cells (green) in the in the proximal tibial growth plate (n=3). The *Axin2Cre*$^{ERT2}$*;R26R*$^{ZsGreen}$ mice received three daily injection of tamoxifen on P23-P25. At P26, the hind limbs were collected. The white dashed lines demarcate the growth plate from the secondary ossification center. Scale bars: 50 µm. Arrows represent *Igf1r*-expressing Axin2+ cells.

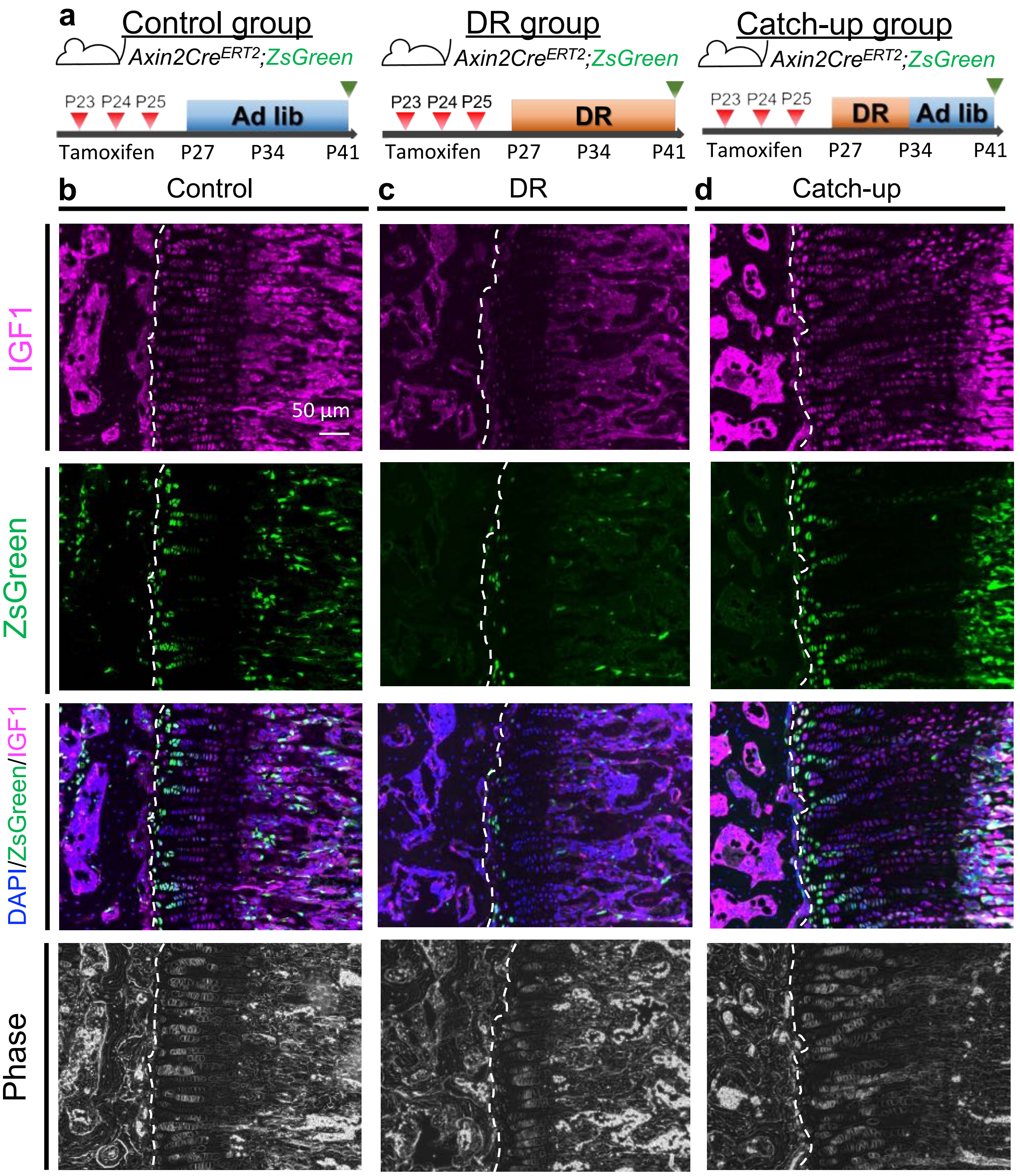


**Supplementary Figure S12. IGF1 expression during catch-up growth in the growth plate. a** Schematic diagrams of the mouse catch-up growth model examining the proximal tibial growth plate of C57/B6J mice. The mice were fed ad libitum (control group), subjected to DR from P27-P41, or ad libitum after seven-day DR from P27 (catch-up group). The hind limbs were collected at P41. **b-d** Representative images of immunostaining of IGF1(magenta) and Axin2+ cells (green) in the proximal tibial growth plate (n=3). The white dashed lines demarcate the growth plate from the secondary ossification center (left). Scale bars: 50 μm.

**Supplementary Table S1.** Upstream regulator analysis of the genes changing from resting to proliferative chondrocytes in the growth plate cartilage of 30-day-old mice using Ingenuity Pathway Analysis.

| Upstream Regulator | Activation z-score | P value |
|---|---|---|
| Lipopolysaccharide | 6.263 | 2.63E-42 |
| Dexamethasone | 1.579 | 7.63E-38 |
| TNF | 5.554 | 6.03E-37 |
| TGFB1 | 3.603 | 2.04E-32 |
| IFNG | 4.349 | 3.07E-29 |
| Beta-estradiol | 1.195 | 1.35E-27 |
| GATA2 | 4.805 | 6.08E-25 |
| Tretinoin | 5.841 | 1.29E-22 |
| Progesterone | -0.108 | 1.72E-22 |
| IL1B | 4.242 | 4.98E-22 |
| AGT | 4.406 | 1.9E-20 |
| TCL1A | 2.813 | 2.65E-19 |
| SPI1 | 4.243 | 4.37E-19 |
| IL6 | 5.108 | 1.48E-18 |
| CSF1 | 1.183 | 9.06E-18 |
| HRAS | 0.178 | 1.47E-17 |
| FGF2 | 2.897 | 3.02E-17 |
| Immunoglobulin | -0.631 | 3.37E-17 |

| | | |
|---|---|---|
| KRAS | -1.065 | 3.89E-17 |
| STAT3 | 2.69 | 6.4E-17 |
| IL4 | 2.254 | 9.64E-17 |
| D-glucose | 1.607 | 4.27E-16 |
| IGF1 | 1.804 | 4.44E-16 |
| NFKBIA | 3.305 | 7.03E-16 |
| ERBB2 | -0.983 | 8.07E-16 |
